# Benchmarking of deep learning algorithms for 3D instance segmentation of confocal image datasets

**DOI:** 10.1101/2021.06.09.447748

**Authors:** Anuradha Kar, Manuel Petit, Yassin Refahi, Guillaume Cerutt, Christophe Godin, Jan Traas

## Abstract

Segmenting three dimensional microscopy images is essential for understanding phenomena like morphogenesis, cell division, cellular growth and genetic expression patterns. Recently, deep learning (DL) pipelines have been developed which claim to provide high accuracy segmentation of cellular images and are increasingly considered as the state-of-the-art for image segmentation problems. However, it remains difficult to define their relative performances as the concurrent diversity and lack of uniform evaluation strategies makes it difficult to know how their results compare. In this paper, we first made an inventory of the available DL methods for 3 dimensional (3D) cell segmentation. We next implemented and quantitatively compared a number of representative DL pipelines, alongside a highly efficient non-DL method named MARS. The DL methods were trained on a common dataset of 3D cellular confocal microscopy images. Their segmentation accuracies were also tested in the presence of different image artifacts. A specific method for segmentation quality evaluation was adopted which isolates segmentation errors due to under/over segmentation. This is complemented with a 3D visualization strategy for interactive exploration of segmentation quality. Our analysis shows that the DL pipelines have different levels of accuracy. Two of them, which are end to end 3D and were originally designed for cell boundary detection, show high performance, and offer clear advantages in terms of adaptability to new data.

**Author summary:** In recent years a number of deep learning (DL) algorithms based on computational neural networks have been developed which claim to achieve high accuracy and automatic segmentation of 3D microscopy images. Although these algorithms have received considerable attention in the literature, it is difficult to evaluate their relative performances, while it remains unclear whether they really perform better than other, more classical segmentation methods.

To clarify these issues, we performed a detailed, quantitative analysis of a number of representative DL pipelines for cell instance segmentation from 3D confocal microscopy image datasets. We developed a protocol for benchmarking the performances of such DL based segmentation pipelines using common training and test datasets, evaluation metrics and visualizations. Using this protocol, we evaluated and compared four different DL pipelines to identify their strengths and limitations. A high performance non-DL method was also included in the evaluation. We show that DL pipelines may show significant differences in their performances depending on their model architecture and pipeline components but overall show excellent adaptability to unseen data. We also show that our benchmarking protocol could be extended to a variety of segmentation pipelines and datasets.

## Introduction

The use of 3 dimensional, quantitative (3D) microscopy has become essential for understanding morphogenesis at cellular resolution, including cell division and growth as well as the regulation of gene expression [1]. In this context, image segmentation to identify individual cells in large datasets is a critical step. Segmentation methods broadly belong to two types, namely ‘semantic segmentation’ in which each pixel within an image is associated with one of the predefined categories of objects present in the image. The other type, which is of interest in this paper, is ‘instance segmentation’ [2]. This type of method goes one step further by associating each pixel with an independent object within the image. Segmenting cells from microscopy images falls within this second type of problem. It involves locating the cell contours and cell interiors such that each cell within the image may be identified as an independent entity [3]. High accuracy cell instance segmentation is essential to capture significant biological and morphological information such as cell volumes, shapes, growth rates and lineages [4].

A number of computational approaches have been developed for instance segmentation (e.g. [1, 5], [6, 7], [8], [9, 10] such as for example the commonly used watershed, graph partitioning and gradient based methods. In watershed approaches seed regions are first detected using criteria like local intensity minima or user provided markers. Starting from the seed locations, these techniques group neighboring pixels by imposing similarity measures until all the individual regions are identified. In graph partitioning, the image is treated as a graph, with the image pixels as its vertices. Subsequently pixels with similar characteristics are clustered into regions also called superpixels. Superpixels represent a group of pixels sharing some common characteristics such as pixel intensity. In some graph based approaches such as [11–13], superpixels are first estimated by oversegmenting an image followed by graph partitioning to aggregate these super-pixels into efficiently segmented regions of the image. Gradient based methods use edge or region descriptors to drive a predefined contour shape (usually rectangles or ellipses) and progressively fit them to accurate object boundaries, based on local intensity gradients [14, 15] .

Common challenges faced by these segmentation methods arise in low contrast images containing fuzzy cell boundaries. This might be due to the presence of nearby tissue structures as well as anisotropy of the microscope that perturb signal quality, poor intensity in deeper cell layers as well as blur and random intensity gradients arising from varied acquisition protocols [16] [17]. Some errors can also be due to the fact that cell wall membrane markers are not homogenous at tissue and organ level: in some regions the cell membrane is very well marked resulting in an intense signal while in the other regions this may not be the case. These different problems lead to segmentation errors such as incorrect cell boundary estimation, single cell regions mistakenly split into multiple regions (over-segmentation), or multiple cell instances fusing to produce a condensed region (under-segmentation).

In recent years, a number of computational approaches based on large neural networks (commonly known as deep learning or DL) [18] have been developed for image segmentation [19] [20, 21]. The key advantages of DL based segmentation algorithms include automatic identification of image features, high segmentation accuracy, requirement of minimum human intervention (after the training phase), no need for manual parameter tuning during prediction and very fast inferential capabilities. These DL algorithms are made of computational units (‘neurons’), which are organized into multiple interconnected layers. For training a network, one needs to provide input training data (e.g images) and the corresponding target output (ground truth). Each network layer transforms the input data from the previous level into a more abstract feature map representation for the next level. The final output of the network is compared with the ground truth using a loss (or cost) function. Learning in a neural network involves repeating this process and automated tuning of the network parameters multiple times. By passing the full set of training data through the DL network a number of times (also termed ‘epochs’) the network estimates the optimal mapping function between the input and the target or ground truth data. The number of epochs can be in the order of hundreds to thousands depending on the type of data and the network. The training will run until the training error is minimized. Thereafter, in a ‘recognition’ phase, the neural network with these learnt parameters can be used to identify patterns in previously unseen data.

In case of application of deep learning for image segmentation, the training and inference process is identical as above. The input data comprises raw images (grayscale or RGB) and the ground truth data are composed of highly precise segmentations of these input images where the desired regions are labeled.

Instance segmentation using deep learning is a challenging task especially for 3D data due to large computational time and memory requirements for extracting individual object instances from 3D data [22] [23] Therefore, the current trend in deep learning based segmentation methods is to proceed in two steps. Firstly deep networks are used to provide high quality semantic segmentation outputs. This involves the extraction of several classes of objects within an image such as cell boundaries, cell interiors and background. These DL outputs are then used with traditional segmentation methods to achieve the final high accuracy and automatic instance segmentation even in images with noise and poor signal quality [24]. A generic workflow of such a deep learning based instance segmentation process is shown in Fig 1.

**Fig 1.**
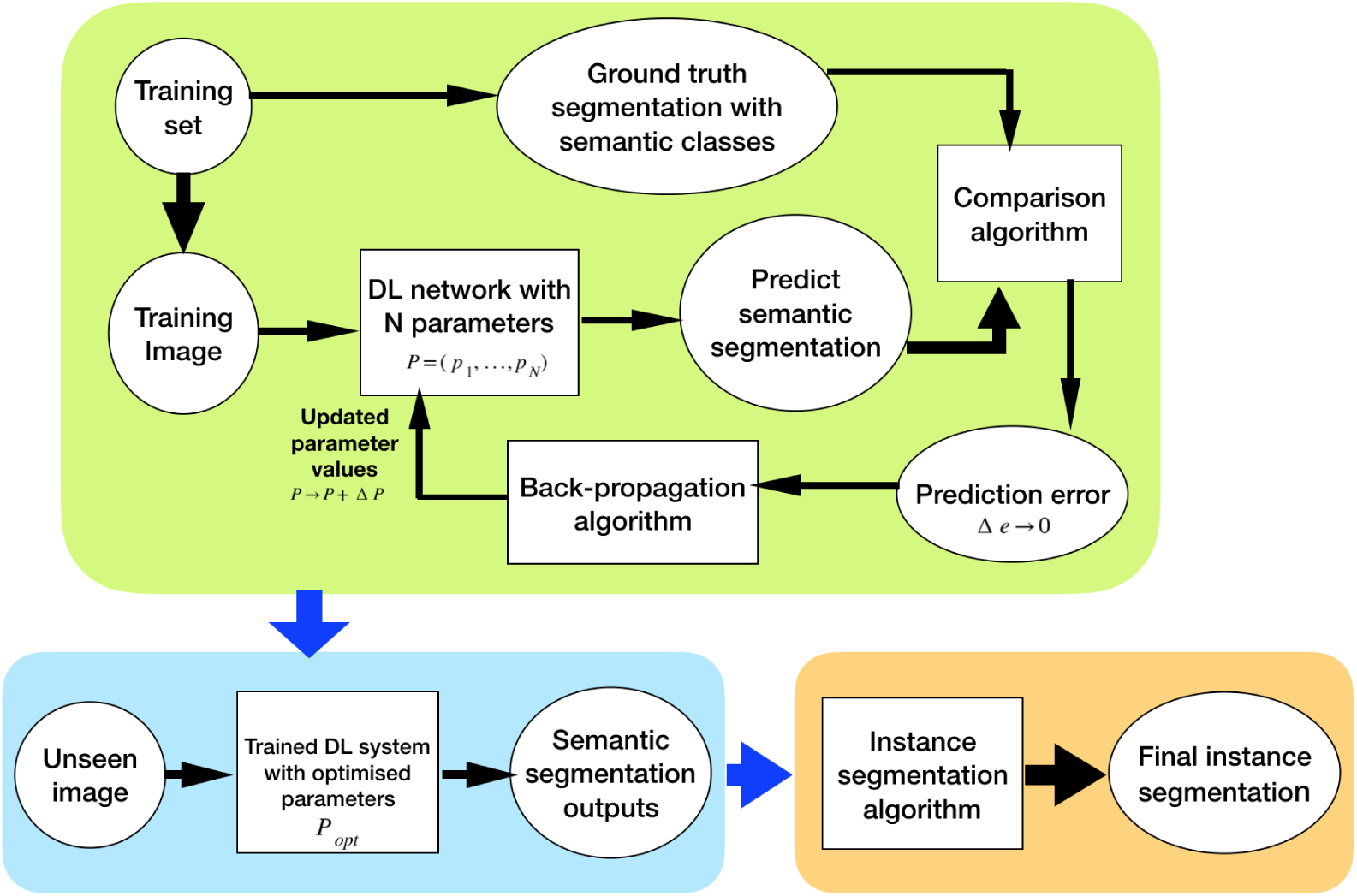
Generic workflow of a deep learning (DL) based image segmentation pipeline. The DL network is first trained to produce a semantic segmentation which corresponds as closely as possible to a given groundtruth. The trained network is then used to segment unseen images. The resulting semantic segmentation is then further processed to obtain the final instance segmentation.

In contemporary deep learning literature, two types of architecture for segmentation are commonly used: the ones based on the UNet/residual UNet network [25] [26] and the approaches using the region proposal networks or RCNNs (Region Based Convolutional Neural Networks) [27]. We will only briefly present both the general properties of both types of networks. The UNet [25] has a symmetric deep learning architecture. One part (called encoder) extracts the image features and the other part (named decoder) combines the features and spatial information to obtain the semantic segmentation, for example cell boundaries, cell body and image background. In order to obtain the instance segmentation, this is followed by methods such as watershed or graph partitioning. Examples of UNet based 2D and 3D segmentation algorithms include e.g. [28] [29] [24].

Besides UNet, the other state-of-the-art deep learning architecture is a Region based convolutional neural network or RCNN such as Mask-RCNN or MRCNN [30]. MRCNN differs from UNet as the former includes modules for object detection and classification unlike the UNets. MRCNN has been used for high accuracy and automatic segmentation of microscopy images in several works(e.g. [31] [32] [33]).

There currently exists a large number of deep learning pipelines (we have identified and reviewed up to 35 works in the last 5 years in Supporting information file S4), where variants of both the above architectures are used to address specific challenges in segmentation such as sparse data sets, availability of partial ground truths, temporal information, etc (see Supporting information file S4 for a more extensive review). However the diversity of the currently available pipelines and inconsistent use of segmentation accuracy metrics makes it difficult to characterize and evaluate their relative performance based on the literature. The presence of such diversity has motivated several benchmarking studies such as [34] and [35]. Ulman et al [34], describe a thorough evaluation of 21 cell tracking pipelines. One of these includes a 3D UNet deep learning system for segmentation. This study evaluates the capability of the methods to segment and track correctly different types of data (optical and fluorescent imaging, single and densely packed cells). Funke et al [35], compare four epithelial cell tracking algorithms using eight time-lapse series of epithelial cell images. Although both papers highlight the importance of accurate image segmentation and underline the performance of DL, the characterisation of the method errors is rather focused on the cell tracking part. In this paper we focus in detail on the segmentation itself, comparing extensively and quantitatively the capacity of a number of selected DL protocols to accurately segment 3D images. To do so, we retrained the DL systems on a common benchmark 3D dataset and analyzed segmentation characteristics of each pipeline at cellular resolution. The pipelines used here are based on either UNet or RCNN architectures and were selected from the literature based on the following criteria. (i) Firstly, as the focus of this work is on 3D confocal datasets the pipelines are built for 3D instance segmentation of static images. The analysis of temporal information or specific architectures for cell or particle tracking are not included as these are extensively covered in [34] and [35] . (ii) Next, the pipeline implementations including pre and postprocessing methods are available in open-source repositories. (iii) To ensure that the pipelines are reproducible properly on other machines, the training dataset used originally by the authors are available publicly. (iv) Lastly, the DL pipelines are trainable with new datasets. Based on these criteria, we identified four pipelines ([24], [27] [29] [36]), which we further describe below.

The first pipeline is an adapted version of Plantseg [29] which can be trained using 3D images composed of voxels. It uses a variant (see Materials and methods section) of 3D UNet called residual 3D-UNet [26] for the prediction of cell boundaries in 3D, resulting in a semantic segmentation. These are then used in a post-processing step for estimating the final instance segmentation using graph partitioning. Examples of graph partitioning include GASP [8], Mutex Watershed [37], Multicut [13].

The second deep learning pipeline [24] comprises a 3D UNet which can be trained using 3D confocal images (i.e. composed of voxels) for prediction of cell boundary, cell interior and image background regions (as 3D images). These semantic outputs of the 3D UNet are then used to generate a seed image for watershed based post-processing. Seeds in watershed segmentation indicate locations within images from where growing of connected regions starts in the image watershed map. The seed images produced from the UNet outputs in this pipeline are therefore used to perform 3D watershed and obtain the final segmentation output.

The third pipeline is adapted from Cellpose [36]. It uses a residual 2D-UNet architecture which should be trained using 2D images (composed of pixels). The 2D trained UNet predicts horizontal (X) and vertical (Y) vector gradients of pixel values, or flows, along with a pixel probability map (indicating whether pixels are inside or outside of the cell regions) for each 2D image. By following the vector fields the pixels corresponding to each cell region are clustered around the cell center. This is how 2D gradients in XY, YZ and ZX planes are estimated. These 6 gradients are averaged together to find 3D vector gradients.These 3D gradients are used to estimate the cell regions in 3D.

The fourth deep learning pipeline is adapted by the authors of this paper from the well documented open Mask RCNN repository [27] and the 3D segmentation concept using this model is inspired from [32]. For the Mask-RCNN based segmentation, a hybrid approach is adopted as shown in Fig 2. The pipeline uses a MRCNN algorithm which is trained using 2D image data to predict which pixels belong to cell areas and which do not in each Z slice of a 3D volume leading to a semantic segmentation. Then the Z slices containing the identified cell regions are stacked into a binary 3D seed image. The cell regions in this binary image are labeled using the connected component approach, where all voxels belonging to a cell are assigned a unique label . These labeled cell regions are used as seeds for watershed based processing to obtain the final 3D instance segmentation.

**Fig 2.**
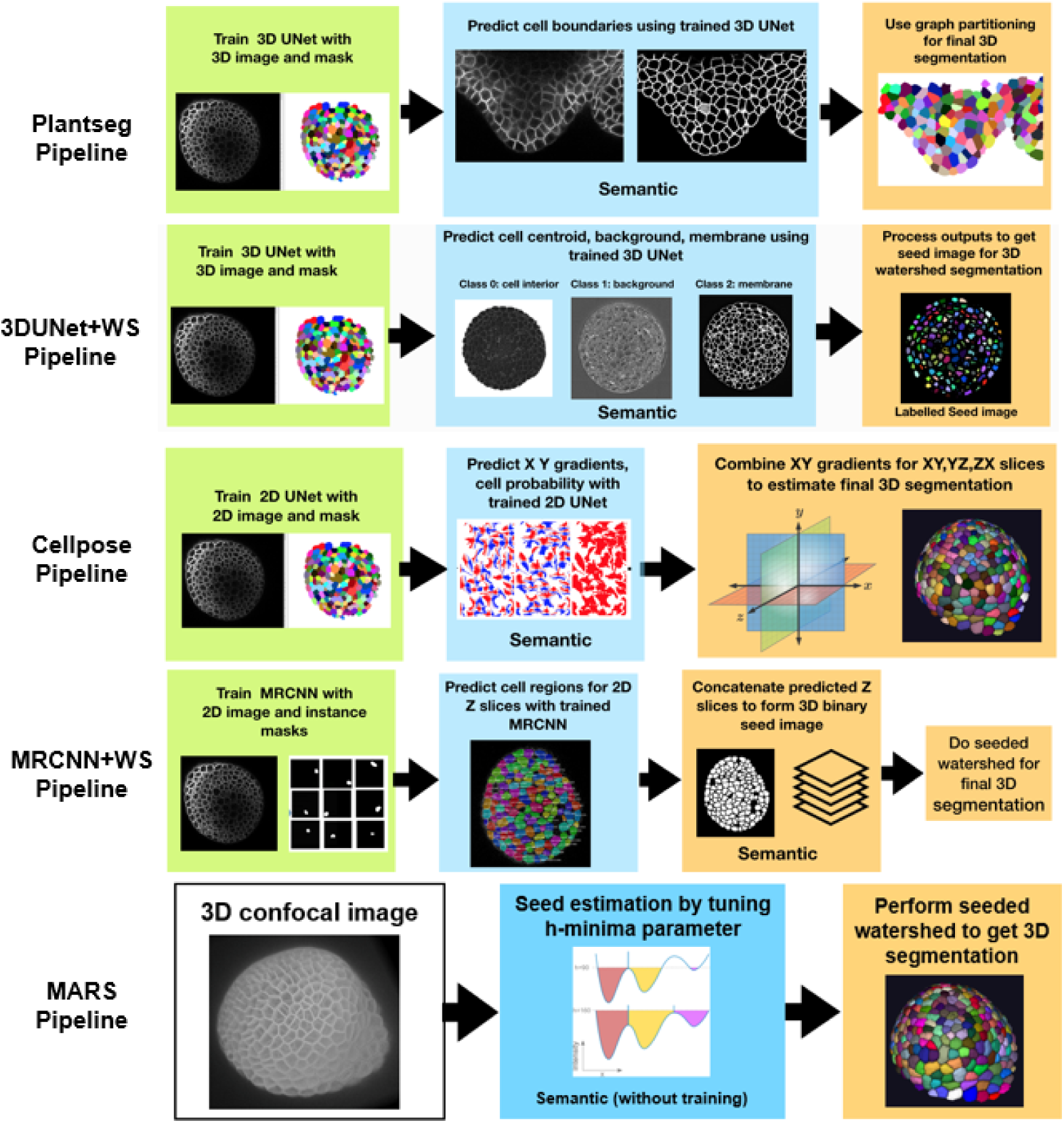
Displaying all the 3D segmentation pipelines together. The green colored boxes indicate the training process for the respective pipeline. The blue boxes indicate the predicted, semantic segmentations generated by the trained DL algorithms, the orange boxes indicate phases of post processing, leading to the final instance segmentation. The MARS pipeline doesn’t include a training or post processing step, but parameter tuning is required.

A further aspect of investigation in this work is to observe how these deep learning pipelines compare to a classical non-deep learning pipeline in terms of segmentation accuracy. They were therefore compared with a watershed based segmentation pipeline named MARS [38], which uses automatic seed detection and watershed segmentation. In the MARS pipeline, the minima of local intensity of the image detected by a h-minima operator is used to initiate seeds in the image which are then used for 3D watershed segmentation of cells. This pipeline therefore does not involve any model training component.

As these five segmentation pipelines have been developed and tested on different datasets and have been characterized using different evaluation metrics, it is difficult to directly compare their performance. For example, in the original papers the Plantseg and UNet+WS pipelines were trained and designed for images having membrane stainings and therefore use a UNet based boundary detection method. The Cellpose model was originally trained with diverse types of bio-images such as those with cytoplasmic and nuclear stains and microscopy images with and without fluorescent membrane markers. The Mask RCNN adopted in this work was originally trained using images from cell nuclei.

Therefore, the first step of the benchmarking protocol was to train the four deep learning pipelines on a common 3D image dataset. We next tested all the 5 segmentation (DL and non-DL) pipelines on a common 3D test image dataset. This was followed by estimating and comparing their performance based on a common set of metrics. Through this protocol we aimed to develop an efficient strategy for quantitative and in depth comparison of any 3D segmentation pipeline that currently exists or is under development. Our results show clear differences in performance between the different pipelines and highlight the adaptability of the DL methods to unseen datasets.

## Results

### A benchmarking protocol for 3D segmentation pipelines

The benchmarking workflow for the segmentation pipelines adapted in this paper is shown in the schematic diagram of Fig 3. All 4 deep learning pipelines were trained following the specifications of respective pipelines (more details in Materials and Methods section) as given in their repositories. For training we used a common set composed of 124 3D original stacks of confocal images from Arabidopsis shoot apical meristems and their ground truth segmentations, which are publicly available, as described in [39]. This is one of the more extensive 3D confocal sets with ground truth publicly available. The trained networks were used to segment two test datasets of floral meristem images (Fig 4A) described in (Refahi et al 2021, see Materials and methods section) and for which ground-truths were available as well. Sample results from these pipelines on one test stack (TS1-00h) are shown in Fig 4B.

**Fig 3.**
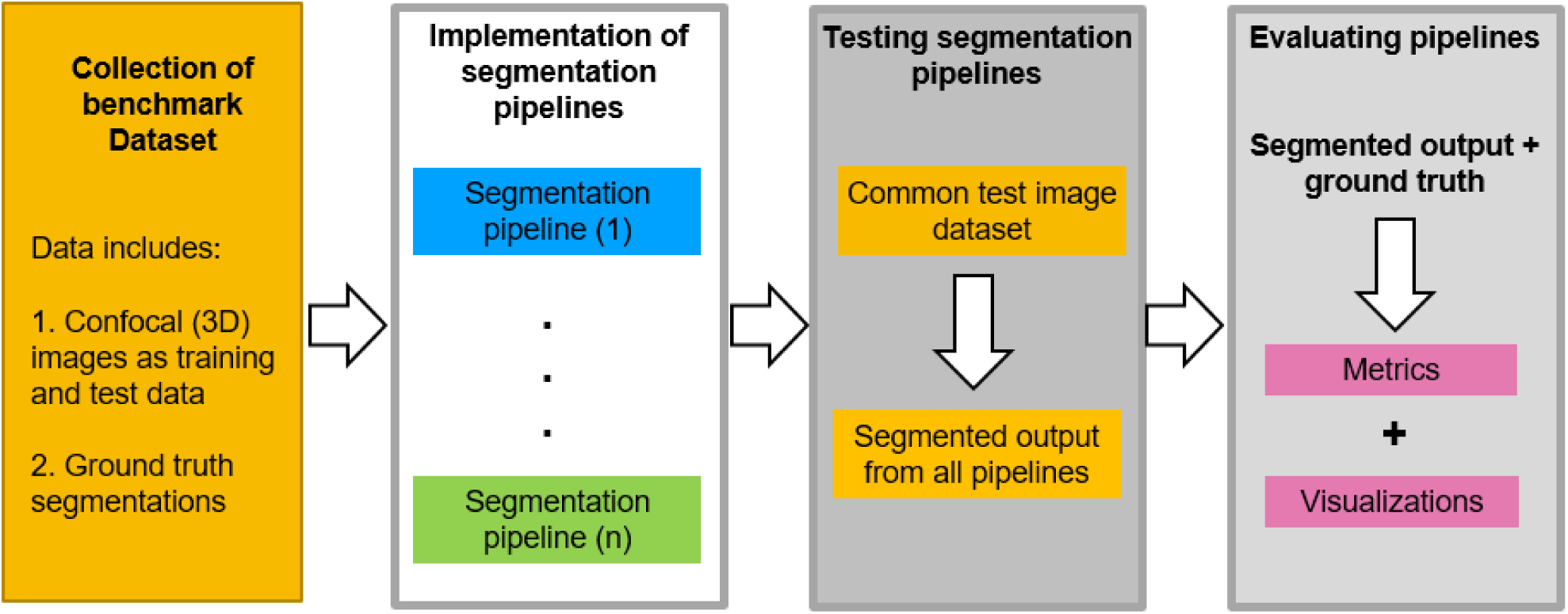
Schematic workflow of the benchmarking process. The evaluation of segmentation pipelines begins with the training of the deep learning models on a common training dataset (confocal images and ground truth). The training and post processing steps for each pipeline are reproduced in the exact way as defined in the respective papers or their repositories. Then the five pipelines are tested on a common test set of images. The test dataset (Fig 4) contains both raw confocal images and their corresponding expert annotated ground truths, and therefore it is possible to assess the segmentation accuracy of the 5 pipelines by comparing segmentation output of each pipeline with the respective ground truth data. Finally the relative accuracy of each method is evaluated using multiple strategies.

**Fig 4.**
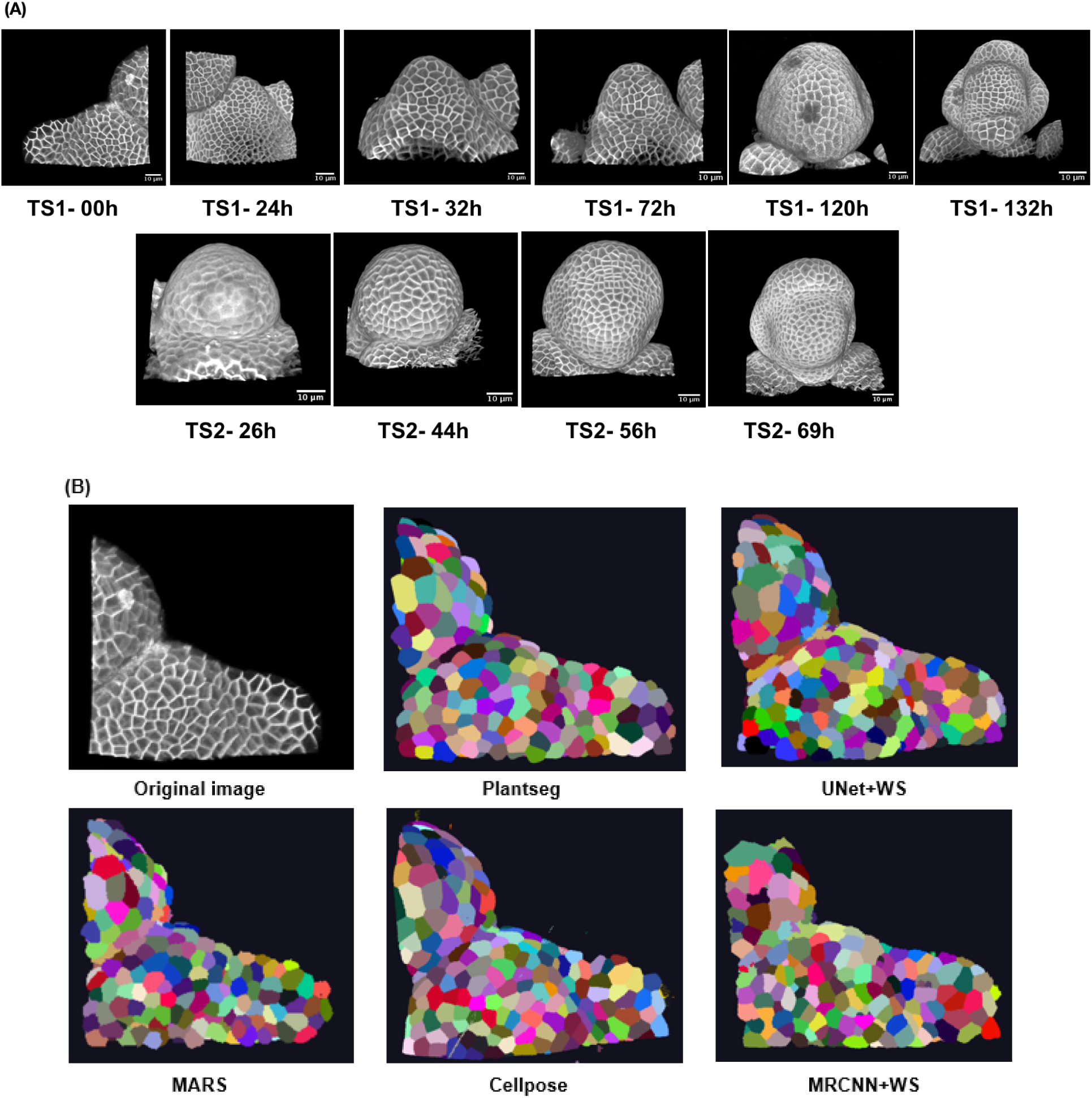
(A) The two test datasets containing a total of 10 confocal image stacks of two different Arabidopsis floral meristems (B) A sample test stack (TS1-00H) and its segmentation by 5 segmentation pipelines.

For MARS, a manual tuning of three parameter values is generally required to obtain optimal segmentation (h-minima, Gaussian smoothing sigma for image and that for seeds, see Materials and Methods section for details on MARS parameters to tune). This can involve many trials before optimal segmentation is obtained and needs expert supervision. For the sake of comparison with the DL methods which were only trained once, we therefore only used one set of parameters. First, the optimal MARS parameters were found for one 3D image and these were then kept constant for the remaining images in the test set.

The other parts of the benchmarking workflow, that is testing and evaluation of the 5 segmentation pipelines through common strategies, metrics and visualizations are detailed below.

### Comparing the segmentation pipelines

We next compared the quality of the segmentations produced by the 5 pipelines. For this purpose we adopted three different strategies (see also Materials and methods for details). First, we analyzed the quality of segmentation on overall stacks, including all cell layers. Next, the confocal stacks were split into different cellular layers (L1, L2, inner) and the segmentation quality of the 5 pipelines for each layer was studied. Finally, we evaluated the segmentation quality on images with commonly occuring (artificially generated) aberrations. In each strategy, several metrics were used for quantitative assessment.

#### Strategy 1: Evaluating segmentation quality for entire image stacks

To estimate the segmentation quality of the outputs from the 5 pipelines, we used their segmented results and the corresponding ground truths of the test datasets. A Volume averaged Jaccard Index (VJI) metric was used to estimate overlap between the predicted segmentations and ground-truths. The VJI used here measures the degree of overlap averaged over the cell volume. In the VJI metric the averaging over cell volume is done to avoid biases arising from the cell sizes on the standard JI. Also metrics that identify the rates of over and under segmentation were applied (details of this metric is in the Materials and Methods section). The rate of over-segmentation is the % of cells in the ground truth associated with multiple regions in the predicted segmentation. Conversely the rate of under-segmentation is the percentage of cases where several regions in the groundtruth are associated with a single cell in the predicted segmentation. Another estimate shown on Table 1 is the percentage of missing cells, which refer to the percentage of cells from the ground truth that are not in the predicted segmentation.

**Table 1:** Mean values (average over the two test datasets) of segmentation evaluation metrics. In MARS the optimal parameter values are only determined for one image (h-minima= 2, Gaussian smoothing sigma for image =0.4).

| **Algorithms** | **Vol. Avg. Jaccard index (VJI)** | **Rate of over-segmentation (%)** | **Rate of under-segmentation (%)** | **Percentage of missing cells (%)** |
| --- | --- | --- | --- | --- |
| **Plantseg** | 0.819 | 4.457 | 6.178 | 2.051 |
| **3D UNet+ WS** | 0.743 | 4.066% | 13.290 | 1.705 |
| **Cellpose** | 0.694 | 13.456 | 10.653 | 4.727 |
| **MaskRCNN+WS** | 0.560 | 14.632 | 22.995 | 9.621 |
| **MARS** | 0.815 | 2.576 | 12.492 | 2.759 |

The results, summarized in Table 1 and Fig 5A-D, reveal a number of differences between the pipelines. Sample results are shown in (Fig 5D)

**Fig 5.**
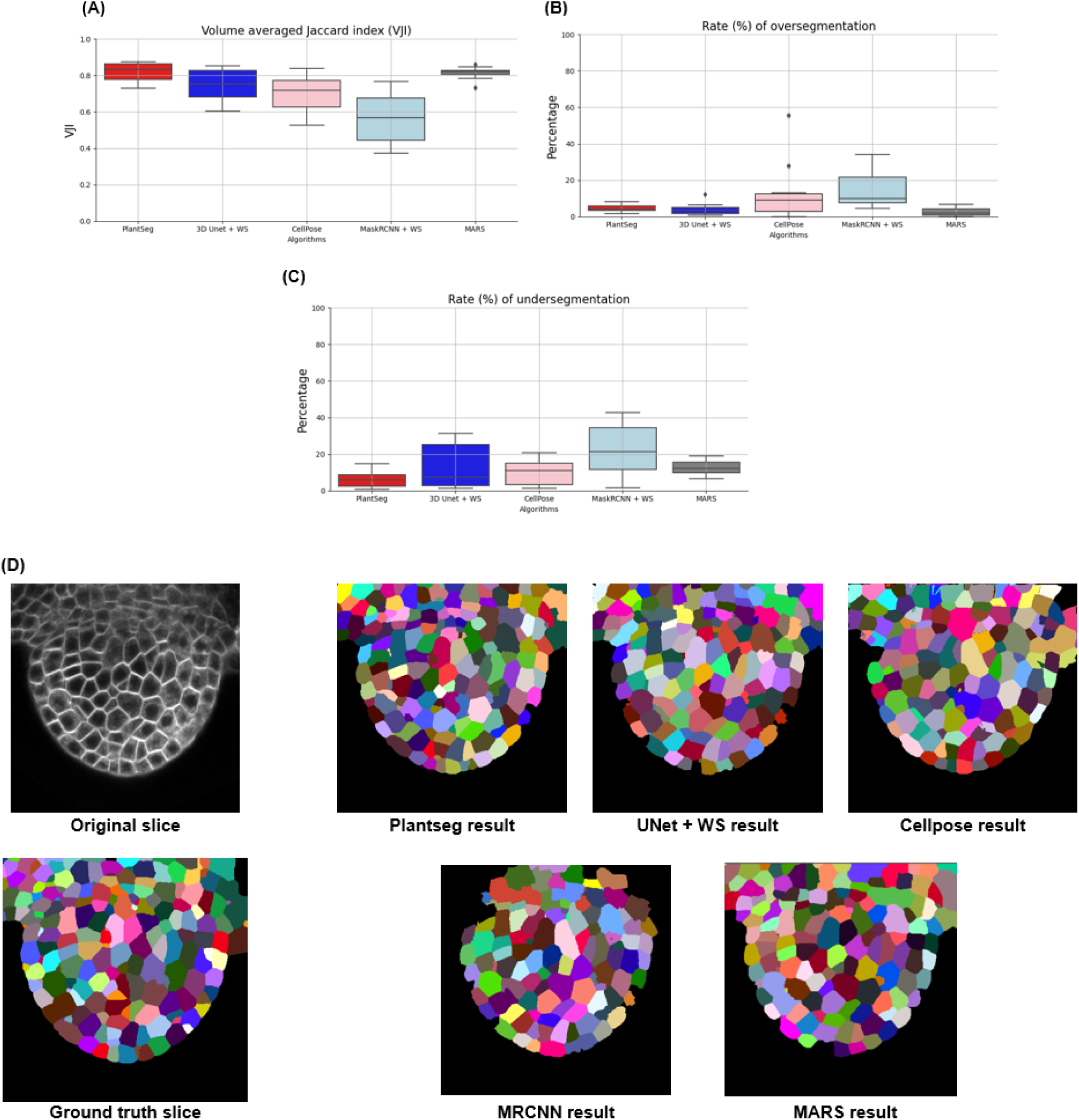
(A) Results of VJI metric from the five segmentation pipelines. Note that VJI is computed for each pair of segmented image/ ground truth image and so the VJI statistics shown above are computed on the values of VJI of the 10 3D test images for each pipeline. (B) and (C) shows rates of over and under segmentation which is computed using a segmented stack and corresponding ground truth stack as input. The distributions shown here are estimated over the results from the two test datasets TS1 and TS2. (D) Example segmentation results by 5 pipelines on a test image slice.

Among all the pipelines, Plantseg performs best as measured using the VJI metric values (Fig 5A), closely followed by MARS and then UNet+Watershed. Cellpose and in particular MRCNN+Watershed perform less well.

With respect to rates of over-segmentation (Fig 5B), The MARS and UNet+Watershed pipelines have lowest rates very closely followed by PlantSeg. The two hybrid pipelines Cellpose and MRCNN+Watershed have higher rates of over-segmentation. The rate of under-segmentation (Fig 5C) is higher for all the pipelines than the rate of over-segmentation, but Plantseg performs better than the others. Overall, we can conclude that the errors in the pipelines are mostly due to under-segmentation.

#### Strategy 2: Segmentation quality evaluation for different cell layers

3D confocal stacks often show different levels of intensity and contrast in different cell depths. In particular in the inner layers, the cell segmentation can be challenging. We therefore tested the performance of the pipelines for their capacity to segment the different cell layers.

For identifying the layers from the ground-truth segmented stacks, we carefully manually classify the cells in contact with the background into three classes : L1, L2 and inner layer. Then an automatic procedure was applied to propagate the classes to the remaining cells. L2 cells were defined as cells in contact with L1 cells but not the background. Inner cells were defined as cells in contact with L2 cells exclusively (Fig 6A).

**Fig 6.**
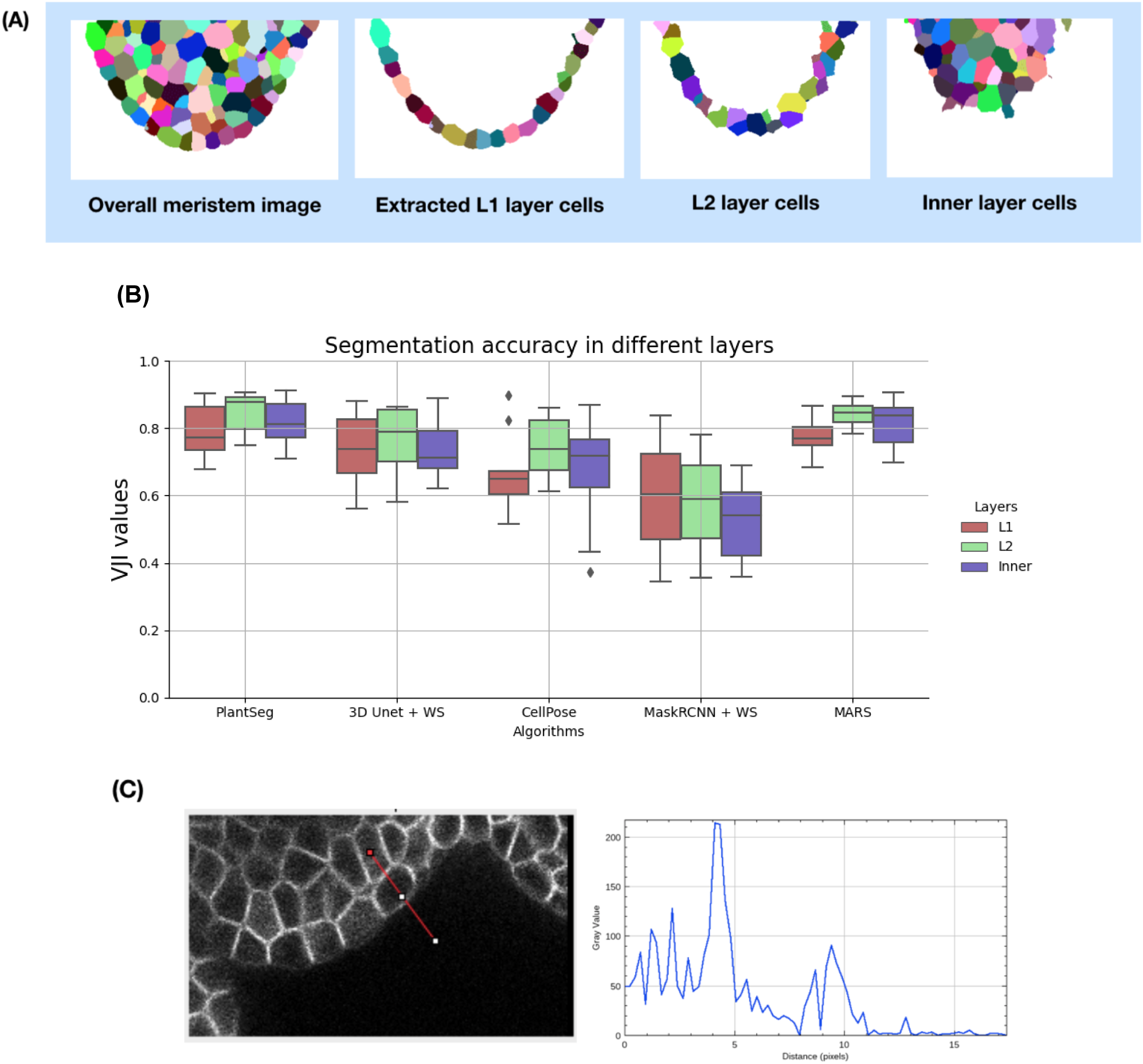
(A) Extracting L1, L2 and inner layers from an input segmented meristem image (B) Estimating segmentation accuracy (VJI) for different cell layers. All stacks from the test dataset are used for this evaluation. (C) Boundary Intensities profile plot for outer and inner layer cells. The gray value at x=0 on the plot on the left is the gray value of the image at the red point of the line segment drawn on the right image.

The variation of segmentation quality of the different pipelines was studied on the basis of the VJI values in the three cellular layers (Fig 6B).

Plantseg and MARS produce the most accurate segmentations in all the layers out of the 5 pipelines and their accuracy doesn’t degrade significantly in the outer or inner layers. The VJI index of the UNet+Watershed pipeline is only slightly lower and nearly the same for all the layers. Cellpose and MRCNN+Watershed perform less well than the others, in all three layers and MRCNN accuracy drops further in the innermost layer. However, with the exception of MaskRCNN, the segmentation of the L2 layer is slightly better than that of the L1. This might be linked to the weak labeling of the outer membranes (Fig 6C) which is often observed.

It may be noted that this layerwise analysis protocol is developed to study how the segmentation pipelines perform when going deeper into the tissue, i.e as the image signal levels get weaker. This analysis is, therefore, also relevant for other tissue types as well, which involves imaging at different cell depths.

#### Strategy 3: Evaluating pipelines on synthetically modified images

Confocal images are often affected by effects such as noise, shadows and motion blur which tend to perturb the image signal. Especially in the inner layers, due to loss of optical signal and scattering, the images contain regions of very poor signal and distortions. In order to study the impact of these variations on the segmentation quality, the effects of noise, blur and intensity variations were simulated on the test set of confocal images. The five segmentation algorithms were then applied to the modified images and the VJI values and rates of under/over segmentations were estimated to observe and compare their robustness against these conditions.

##### Effect of image noise

the electronic detector or amplifier of the imaging equipment can generate noise, which can be modeled using Gaussian statistics [40]. An image to which two different Gaussian noise levels corresponding to noise variances of 0.04 and 0.08 are added is shown in Fig 7A. At higher variance, the noise effect is increased as seen in the PSNR values.

**Fig 7.**
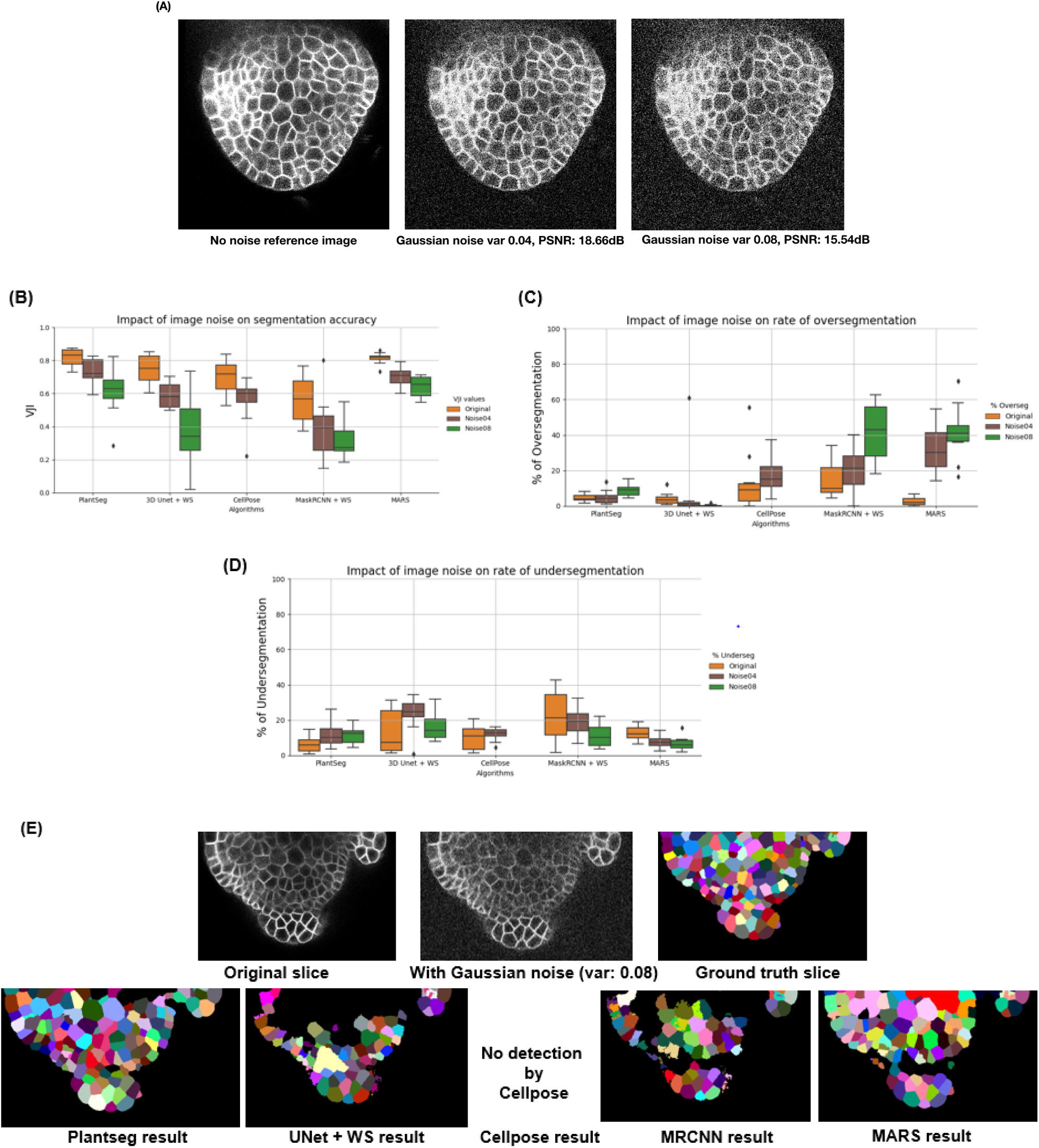
(A) A test image after applying Gaussian noise (var 0.04, 0.08) (B) Variation of segmentation accuracy (VJI) with 3 Gaussian noise variances (C) variation in rates of over-segmentation (D) variation in rates of under-segmentation. Note that for noise variance of 0.08, Cellpose is unable to identify cells. (E) Example results from the five pipelines under the impact of image noise (Gaussian noise variance 0.08)

The five pipelines behaved very differently under the impact of image noise as shown in Fig 7B. In particular UNet+WS, CellPose and MRCNN are very sensitive to Gaussian noise as their accuracy drops sharply when Gaussian noise variance is increased. At a noise variance of 0.08, Cellpose shows no detection. For MRCNN, higher noise leads to loss in identified cell regions, which results in large blob-like regions after watershed based post-processing leading to higher under-segmentation (Fig 7D). The difference with Plantseg could be due to differences in the instance segmentation components. Plantseg uses graph partitioning while UNet+WS and MRCNN pipelines use 3D watershed although the seed identification criteria are different for them. MARS is most sensitive when it comes to oversegmentation (Fig 7C), while PlantSeg is more sensitive in terms of under-segmentation

##### Effect of image blur

Blurring is common in images and can for example be caused by lens aberrations or optical diffraction in the imaging setup [41] . It can also be caused by motion of the objects in the microscope. To simulate blur, the test confocal image is convolved with a horizontal motion blur kernel (material and methods). A sample image before and after blurring is shown in Fig 8A.

**Fig 8.**
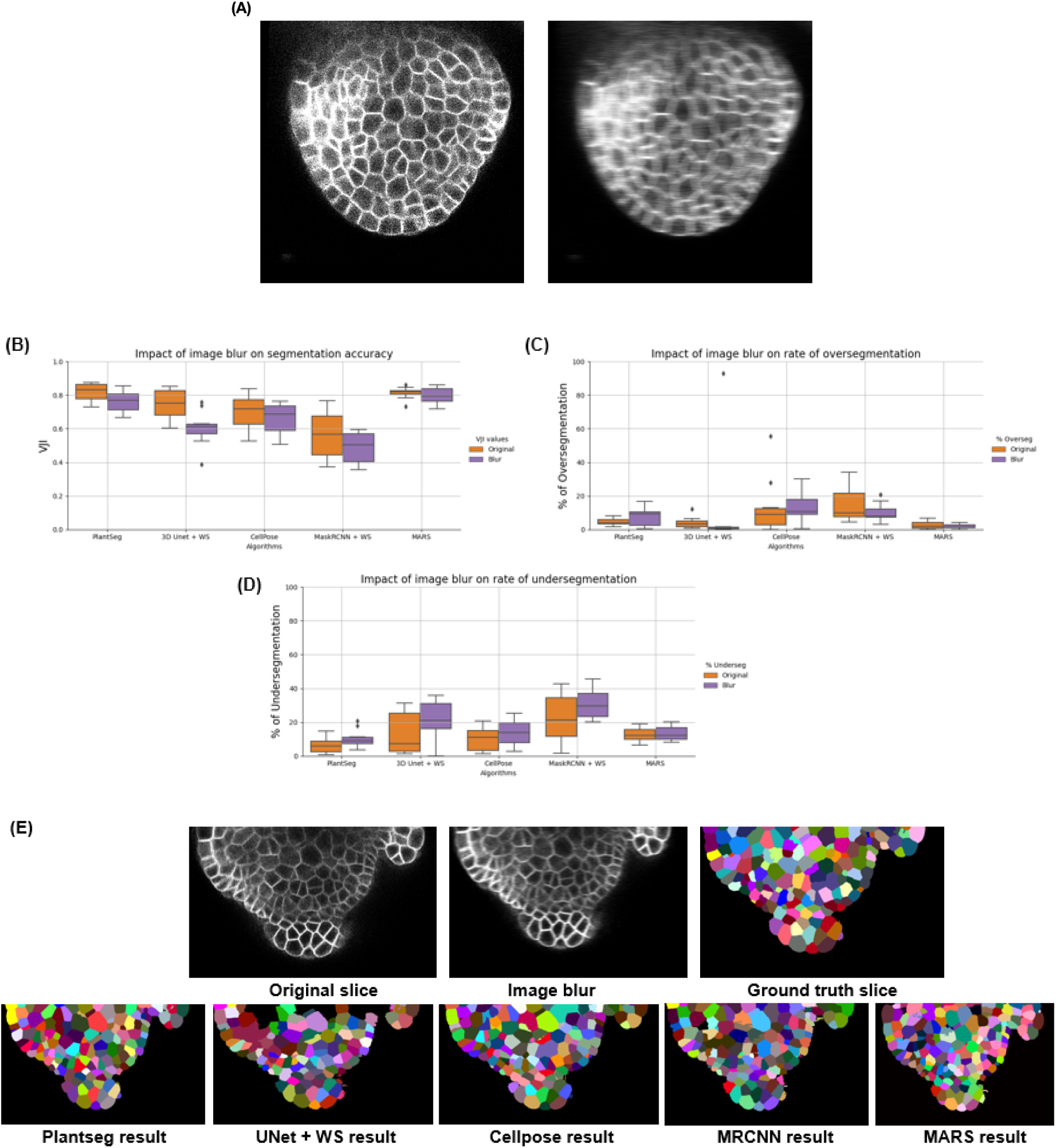
(A) Effect of blurring on an image (B) Comparing segmentation accuracies of pipelines under the effect of image blur (C) Comparing rates of over-segmentation (D) Under-segmentations due to image blur. (E) Results from the five pipelines under the impact of image blur

The 10 test stacks were subjected to the blurring function and were segmented using the 5 pipelines (Fig 8E). VJI values were then computed for each of the results and plotted (Fig 8B). It is seen that the Plantseg, MARS and CellPose pipelines are relatively less affected by blurring, whereas this effect produces larger variability in results in the UNet +WS pipeline. The effects on rates of under-segmentations remain relatively low (Fig 8C). Apart from Plantseg and MARS, the other pipelines suffer higher rates of under-segmentation (Fig 8D) under the impacts of blurring.

##### Image intensity variations

Partially bright regions in microscopy images may be caused by inhomogeneous illumination sources and shadow effects are mostly caused by presence of light absorbing objects or obstructions [42, 43] or due to a non-homogenous cell membrane marker. To emulate the effect of such intensity variations within an image, partial overexposure or underexposure regions (Fig 9A) were imposed on the test images, which were then segmented using the 5 pipelines. The VJI values and rates of over- and under-segmentations for all the results were computed and are shown in the box plots for all stacks in Fig 9B-D. Sample results from all pipelines for over-exposure are shown in Fig 10A and for under-exposure in Fig 10B.

**Fig 9.**
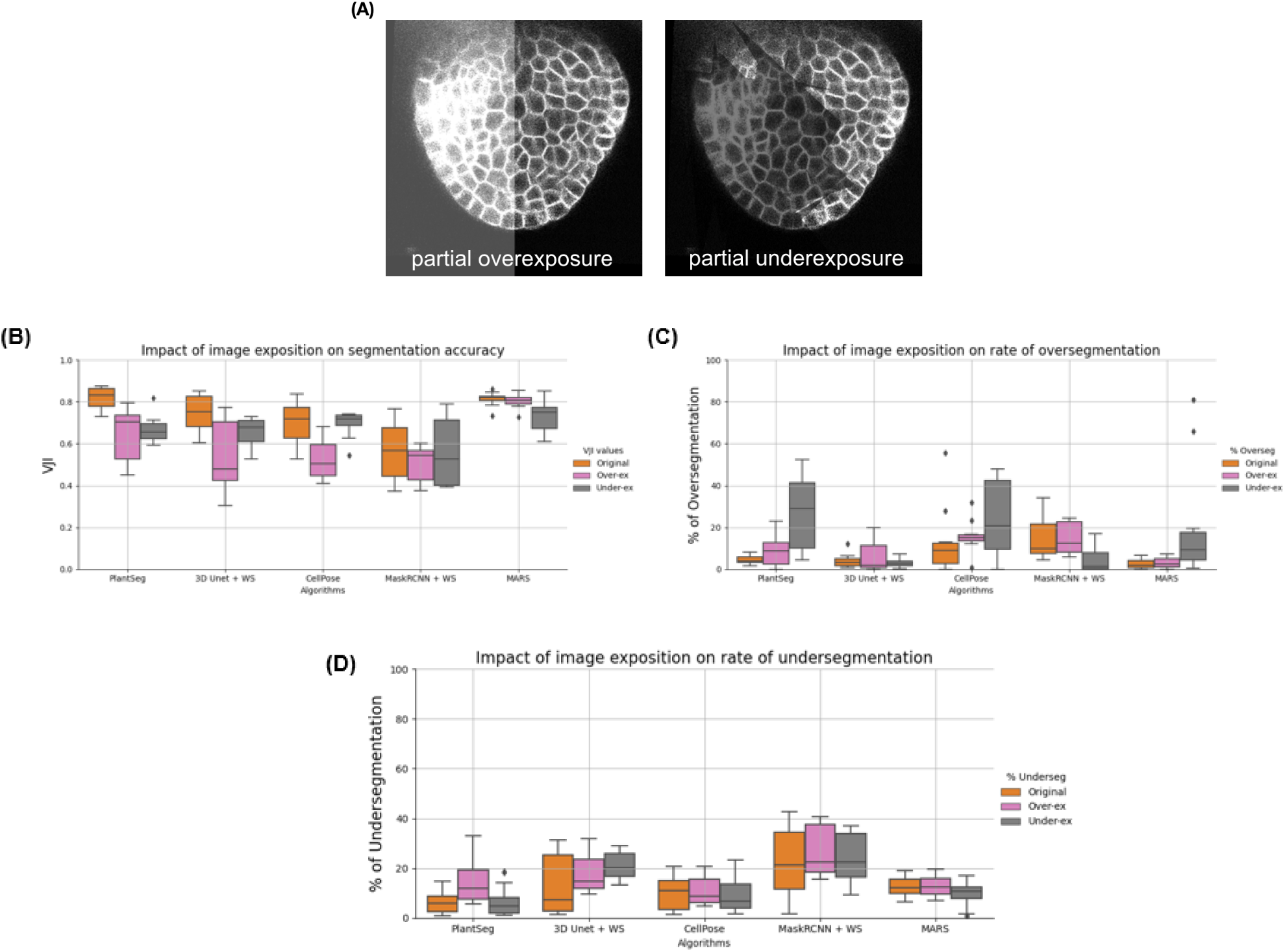
Impact of image exposure levels on segmentation quality of 5 pipelines. (A) examples of partial over and underexposure. In (B) the VJI values for over and under-exposure are plotted together with the original VJI values for unmodified stacks. Similarly in (C) and (D) the rates of over and under-segmentation are plotted for the impacts of over and underexposure alongside those for the unmodified stacks.

**Fig 10.**
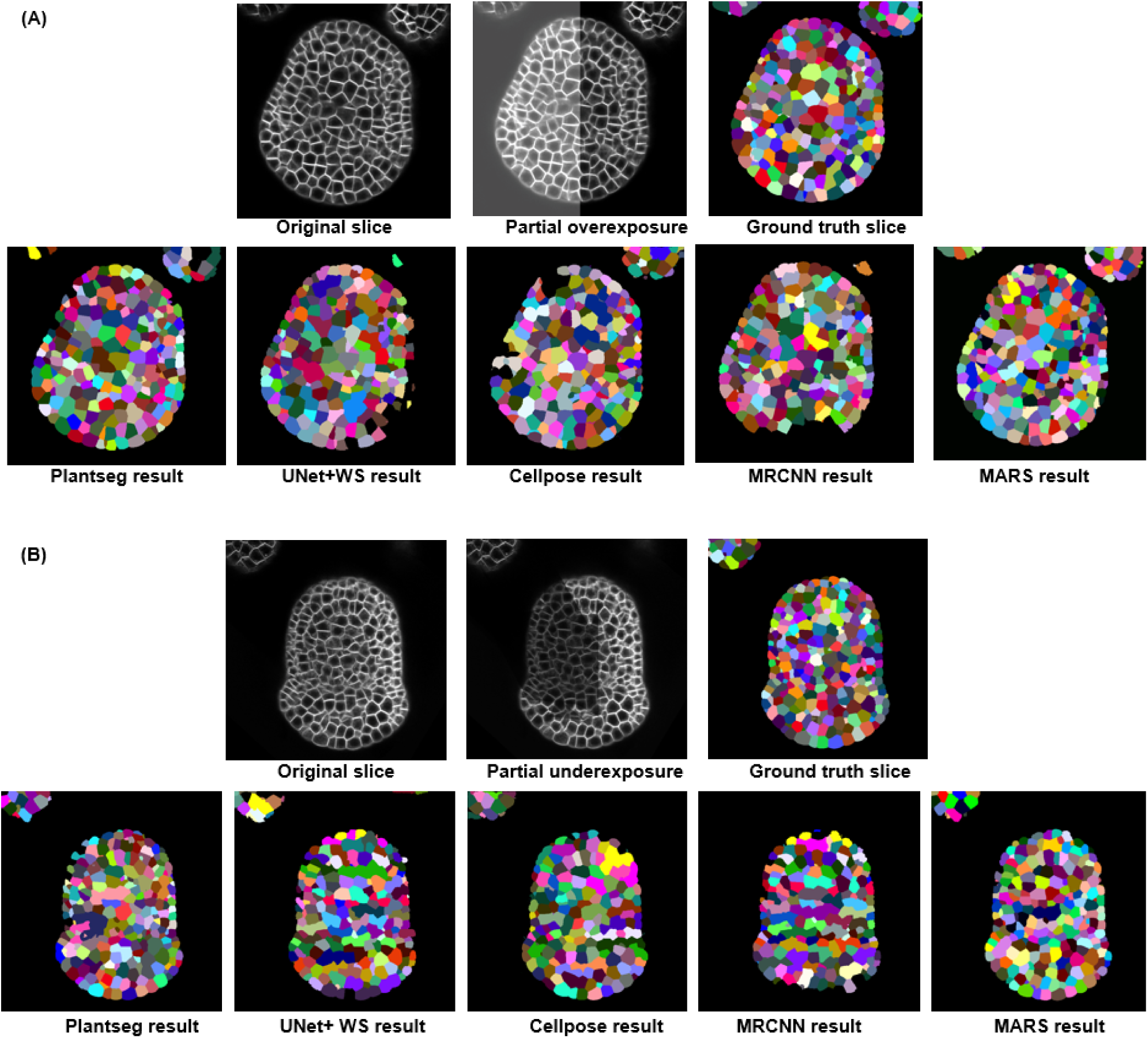
(A) Sample results from the five pipelines under the impact of image over-exposure (B) Results from the five pipelines under the impact of partial underexposure

***Overexposure*** had a strong negative impact on the VJI of Plantseg, UNet+Watershed and Cellpose. The MARS and MaskRCNN pipelines were not affected appreciably. Over-segmentation increased significantly in the Plantseg and Cellpose (Fig 9C) more than other pipelines. Under-segmentation (Fig 9D) was higher in PlantSeg and UNet+WS, while for others it remained at similar levels as on original images.

***Underexposure*** was strongly reflected in the VJI results from Plantseg and also on MARS(Fig 9D). Partial underexposure induced a high degree of over-segmentation in MARS, PlantSeg and CellPose and high rates of under-segmentation in the UNet+WS pipeline. The MaskRCNN based pipeline, although having low overall accuracy, was found to be less sensitive to underexposure.

In conclusion, overall for the Plantseg, UNet+Watershed and Cellpose (or the UNet based) pipelines the effect of image intensity variations appears to be much stronger than image noise or blur effects. MaskRCNN and, to a lesser extent MARS on the other hand, were relatively stable. MARS out of all the pipelines is found to be the most stable under the effect of image artifacts, although it leads to increased over-segmentation in partially underexposed samples.

#### Strategy 4: Evaluating pipelines on unseen data types

We tested the performance of our trained deep learning pipelines and MARS on data different from confocal images of floral meristems, for example on data from other microscopes and tissue types to observe the adaptability of our methods to new and unseen data (the deep learning pipelines were originally trained on shoot apical meristem images). Images from two datasets were used for this and the results from all the pipelines are presented below.

##### a) Ascidian Phalusia mamaliata (PM) embryo images

The 5 pipelines were used for segmenting 3 images from the PM embryo dataset, described in [44]. These were captured using multi-view Light-sheet (MuVi-SPIM) microscopes from fluorescently labeled cell membranes. Ground truth segmentations for these images were also provided in the dataset.

Three test images and their ground truths were taken from each of the above datasets and all five pipelines are used to segment this data. Then the volume averaged Jaccard Index metric was used to estimate the segmentation quality as done with the floral meristem test dataset (Fig 11D).

**Fig 11.**
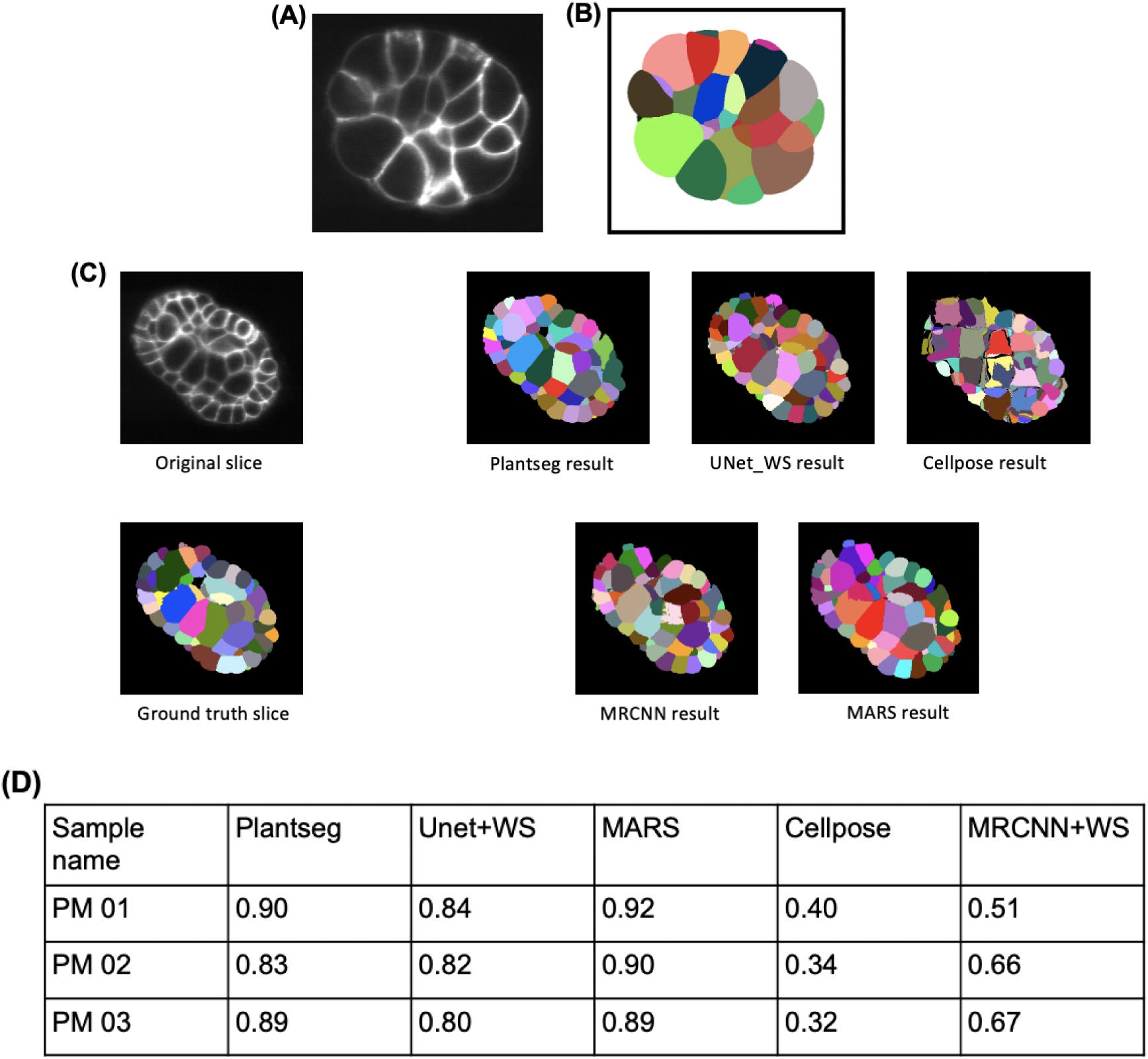
Slice view of a sample (A) Ascidian embryo image and its (B) ground truth segmentation (C) Ascidian embryo image (PM03), ground truth and segmentations by 5 pipelines (D) VJI values for segmentation results using Ascidian PM data and 5 pipelines

The deep learning pipelines Plantseg, UNet+WS provide high accuracy results on completely unseen data without requirement of re-training (Fig 11D). The MARS algorithm after slight tuning of the parameters (mainly image smoothing sigma) for this dataset provides the highest accuracy segmentations. The MRCNN+WS results are similar to what we got previously from this pipeline on floral meristem data. The Cellpose accuracy however falls from their average values observed for floral meristem data.

##### b) Arabidopsis ovule images

3D confocal Images of Arabidopsis thaliana ovules at various developmental stages were taken from the dataset provided by the authors of the Plantseg pipeline [29]. Three stacks from this dataset along with their ground truths were used for evaluating the 5 pipelines. Structurally these are quite different from Arabidopsis floral meristems and the segmentation results from the five pipelines along with the original and ground truth images are shown in Fig 12A. Results from the evaluation are in Fig 12B. Note that MARS needed extensive re-tuning for this dataset (discussed further in the Materials and methods section).

**Fig 12.**
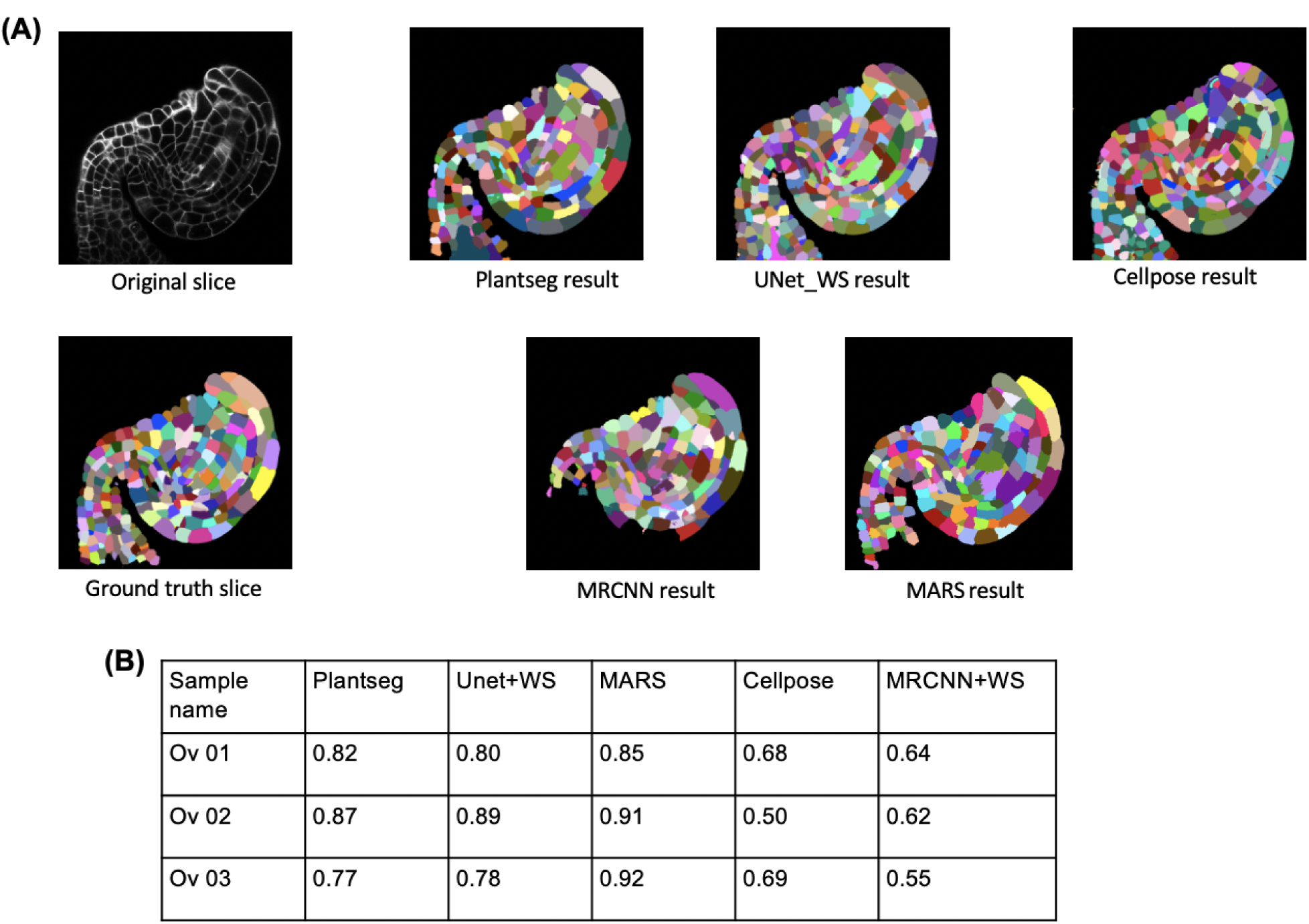
(A) Ovule image and ground truth along with segmentations by 5 pipelines (B) VJI values for segmentation results using ovule data and 5 pipelines

### 3D visualization of segmentation quality

At present, visualization and exploration of 3D image data is possible by software packages such as ImageJ, Paraview or MorphographX [45]. However, only a few tools allow users to project any extrinsic property over an image and to interact with a 3D image dataset at cellular resolution. One such platform is Morphonet [46] (see Supporting information file S2) which is an open source and web-based platform for interactive visualization of 3D morphodynamic datasets. These datasets are created by converting image datasets (3D stacks) into corresponding 3D meshes. So far, Morphonet has been used to project a variety of genetic and morphological information such as gene expression patterns, cell growth rates and anisotropy values on plant and animal tissue images.

Here we have developed a MophoNet based method for interactive 3D visualization of segmentation quality measures, by projecting VJI values on 3D meshes created from 3D confocal images (Fig 13). The 3D meshes were created by the marching cubes algorithm [47] and uploaded on MorphoNet. The VJI values for the image were computed and uploaded as a quantitative property of each individual 3D cell of the image for each segmentation pipeline. The VJI values were then viewed using colormaps on the Morphonet browser.

**Fig 13.**
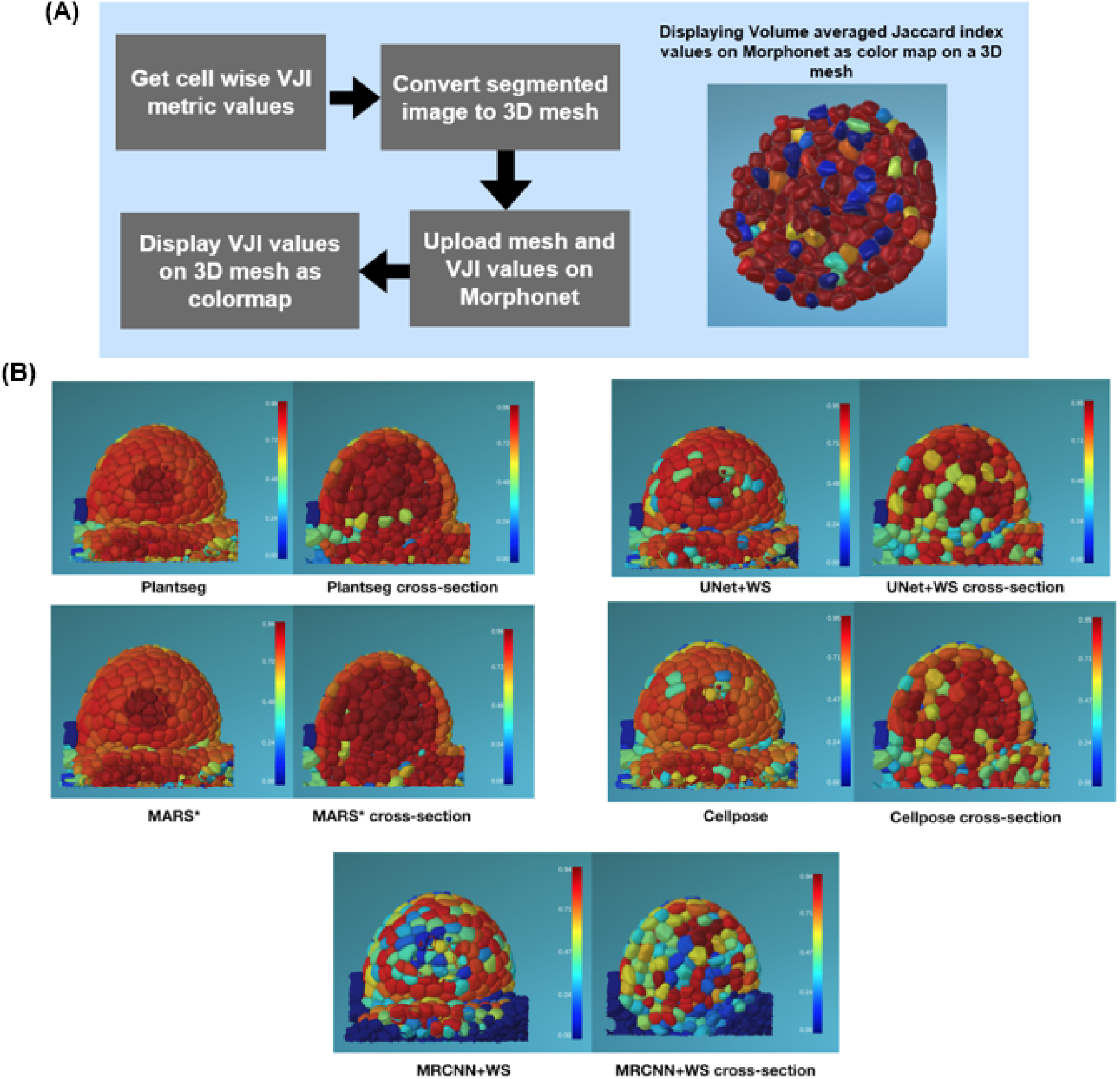
(A) Process to view segmentation quality in 3D on Morphonet.Segmentation quality results (VJI values) for a test stack (TS2-26h) from 5 pipelines displayed on Morphonet. Users can slice through each 3D stack in XYZ directions and check the property (here VJI values) for each cell in the interior layers of the tissue structure. For example, for each pipeline in the above figure the left image shows the full 3D stack and the right image shows the cross section of the same stack after slicing 50% in the Z direction. VJI values are projected as a “property” or colormap on the cells. In this figure a “jet” colormap is used where red represents high and blue represents low VJI values as shown in the color bars alongside.

The steps of the Morphonet based visualization pipeline are provided as open resources in the SegCompare Gitlab repository (described in Supporting information file S1) so that users can upload their own segmentation results on our datasets from any segmentation method or pipeline and benchmark study their accuracy characteristics. The only requirement is that these segmentations should be 16bit labeled .tif files and must have their ground truth segmentations for evaluation. With segmented and ground truth .tif stacks, the steps for Morphonet based segmentation quality visualization includes (see Fig 13A): (a) estimation of cell-by-cell VJI and converting this information to Morphonet compatible format (b) mesh calculation from the ground truth segmented image and converting it to Morphonet compatible .obj format and finally (c) logging in to Morphonet, creating a dataset and uploading the mesh and VJI information. All these steps are documented and implemented in an easy to use Python notebook as described in Supporting information files S1 and S3.

The benefits of the Morphonet based visualization is that a user can: (a) check the VJI value for each cell by clicking on it (b) impose a color mapping of the Jaccard index values so that users can at a glance observe the overall distribution of the VJI values on the cells of the 3D stack (Fig 13B) as well as locate cell regions where poor Jaccard values are found or concentrated (c) interact with segmented data in 3D, e.g rotate 360 degrees or zoom into cells or slice through the segmented data in XYZ directions and inspect segmentation quality in any inner cell layers (d) requires no special software or hardware installation. Also data uploaded on Morphonet can be shared with multiple users who can directly access it on the Morphonet platform. Examples of viewing segmentation quality on Morphonet is illustrated in Fig 13B where mesh belonging to a test stack is uploaded on Morphonet and the VJI values for all the segmentation pipelines are projected on it individually to observe the cell-by-cell variation in segmentation quality.

Sample segmentation accuracy data and 3D meshes in Morphonet compatible format are provided in our repository : https://figshare.com/projects/3D_segmentation_and_evaluation/101120

Sample videos demonstrating various aspects of the 3D visualization process on Morphonet may also be found under “Videos” in our repository described in Supporting information file S3: https://doi.org/10.6084/m9.figshare.14686872.v1

## Discussion

In this study we developed a benchmarking procedure for evaluating 3D segmentation pipelines and used it to analyze 4 deep learning segmentation pipelines and a non-deep learning one. Initially it was difficult to predict the relative performance of the individual pipelines, because they were applied on diverse datasets and used different evaluation metrics. Most common evaluation strategies consist of estimating simple numerical metric values, which also vary widely between the methods. For example in the Plantseg original paper [29], the variation of information metric is used, while other studies are based on the Jaccard index, such as the SEG score in [34], [35] or the aggregated Jaccard index metric (AJI) in the original UNet+WS paper (24). The numerical results from these different metrics are difficult to compare since they are based on different concepts and/or use different datasets. We have used here the volume averaged Jaccard Index. Like the SEG score [34, 35] it is an average Jaccard Index of all pairs of compared regions, but takes into account the cell volume, thus reflecting the error per volume, rather than per cell. Therefore the VJI is more adapted to our data, where cell size can vary considerably.

Another problem is that the simple segmentation accuracy estimates do not provide a detailed view about the types of segmentation errors present in the segmentation results or how the errors are spatially distributed. Therefore, in addition to the VJI metric we evaluated the segmentations by measuring the rates of over- and under-segmentations which indicates the strengths and weaknesses of specific segmentation procedures. This spatial distribution of segmentation quality was further addressed through a layerwise evaluation in this paper. The range of metrics presented here provided an in depth analysis of the type and magnitude of the errors as well as their spatial distributions produced. Each of the metrics gives its own specific information which finally allowed us to define the advantages and shortcomings of each segmentation pipeline. We have also provided full details and coding implementations of the metrics used in the benchmarking process so that they can be re-used by others.

The MorphoNet based interactive evaluation is a fast and efficient way to visualize segmentation quality superimposed on 3D representations. It comprises 3 steps - calculation of cell by cell VJI values for a pair of stacks, creating a 3D mesh and uploading these on Morphonet using their Python API (Supporting information file S2). All these steps are covered in two Python notebooks provided in the SegCompare Gitlab repository (Supporting information file S1). With this visualization, users can navigate through 3D objects in an image and study the segmentation quality for different 3D sections by slicing through the image data (converted to 3D meshes). Since the Morphonet visualization is browser based, it does not require any software installation or special hardware for 3D data visualization. This technique may therefore be added as a final step to any segmentation pipeline, deep learning or non deep learning, for a one-step analysis of the 3D segmentation performance.

We also tried to evaluate how much better the deep learning algorithms performed compared to the non-deep learning algorithms which is a recurrent question in deep learning and computer vision research. Results of PlantSeg, the best performing deep learning pipeline of those tested here, were matched by those of the non-deep learning MARS algorithm. However, without tuning, the MARS accuracy may degrade significantly over completely different datasets as observed with the segmentations of ovule images (see Materials and methods section). By contrast, deep learning models, especially PlantSeg, once trained, performed well throughout. With regard to the time for execution, the MARS segmentation would take much longer than the PlantSeg pipeline (25-30 min for an individual stack vs 8 minutes respectively, see Material and methods section).

The quantitative impact of image artifacts on the accuracies of segmentation pipelines are not always analyzed in literature. This is why one of our protocol strategies was to estimate the impact of image artifacts like blur, over and under exposure and 3 levels of Gaussian noise on the VJI values and rates of over and under-segmentation of the 5 segmentation pipelines. The size of our test dataset therefore consisted of a total of 60 3D stacks with 10 original and 50 synthetically modified ones. In addition, completely unseen 3D images from plant and animal datasets were also segmented using our pipelines.

An important question is why the pipelines have different levels of performance. This could be attributed firstly to the construction of the pipelines. We observed that although 2D segmentation pipelines may be adapted to perform 3D segmentations (e.g. as done in Cellpose and MRCNN+WS), the end to end direct 3D segmentation pipelines (such as Plantseg, UNet+WS pipelines) achieved higher accuracies. This trend is observed in the segmentation accuracy results of the pipelines for 10 original stacks as well as with the stacks having artifacts. Another possible reason behind the difference in performance, especially between the two end-to-end 3D pipelines (Plantseg and 3DUNet+WS). could be arising out of the post processing strategies used in them. Plantseg provides two semantic output classes (background and boundaries) while 3DUNet+WS produces three classes (background, boundary and cell interiors). For the final instance segmentation, Plantseg relies on graph partitioning while UNet+WS uses watershed. It may also be noted that the two pipelines Plantseg and UNet+WS are designed specifically for boundary detection of fluorescently labeled cell boundaries which matches the type of data that we have used in our work. Cellpose on the other hand is built for segmenting a wide range of bio-image data types such as cytoplasm, membrane and cells without fluorescent markers. While the UNets of Plantseg and UNet+WS pipeline extracts and operates on the cell boundary images, the Cellpose pipeline involves prediction of spatial gradient features and a cell probability map that determines the inside and outside regions of cells. The Mask RCNN does not operate on the boundary prediction principle at all but uses object detection and classification for prediction of segmented regions. Thus the difference in performance of the four deep learning algorithms could be attributed to their deep learning model architectures, the way the full segmentation pipeline is constructed as well as on the type of data they are applied to.

We also observed that responses of deep learning pipelines are different to training data augmentation (see Supporting information file S5). The performance of the Plantseg pipeline improves on adding images with artifacts (over and underexposed images) in the training set. By contrast, the performance of the UNet+WS pipeline does not generalize well after using the overexposure augmentation as its performance degrades on normal images. Its performance however improves overall with the underexposure augmentation. We therefore found that same augmentations may not necessarily yield better results for all deep learning based segmentation pipelines. It would also be of interest to investigate the performance of these deep learning models if different types of image data (e.g membrane stained, nuclei stained images, images from other tissue types etc) are mixed in the benchmark training set.

### Data and code availability

All the data and coding implementations from this work are available as open resources. The methods for reproduction of the segmentation pipelines and the segmentation evaluation techniques are documented in the Gitlab repository named SegCompare (https://gitlab.inria.fr/mosaic/publications/seg_compare) along with relevant resources in Jupyter notebooks. The 3D confocal training and test datasets used in this work are provided in open data repositories. These are described in detail in Supporting information files S1 and S3.

## Materials and Methods

### Training data

The training data for all the deep learning algorithms comprise 3D confocal image stacks and their corresponding ground truth segmentations. The structure of a 3D confocal image is shown in Fig 14.

**Fig 14.**
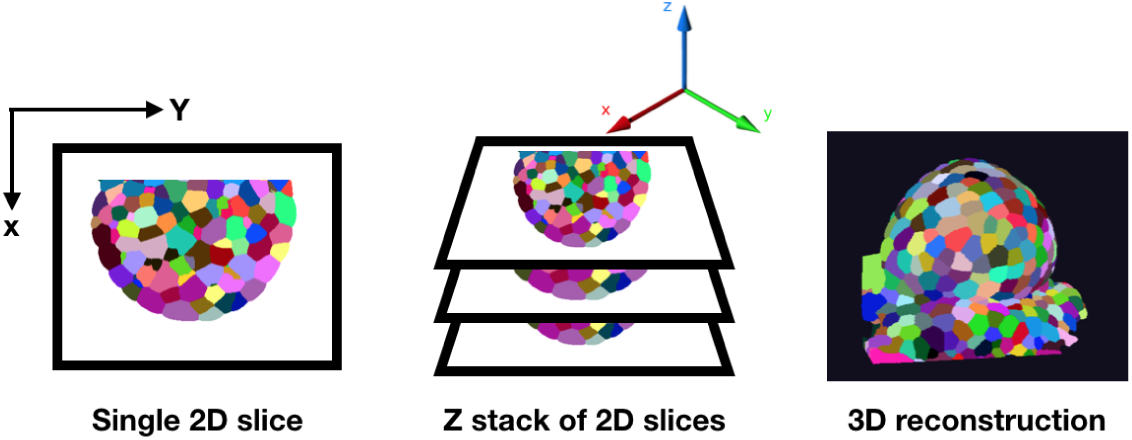
A confocal image is made up by scanning through each point on a 2D plane of an object. The 3D confocal image is made up of such 2D frames stacked along the Z axis. Using the 2D Z slices a full 3D view of the object can be reconstructed.

The source of the training data is Arabidopsis thaliana shoot apical meristems (SAM) which is a multicellular tissue (Fig 15A, C) where cells at the surface grow radially outward from the central to the peripheral zone. This data has been previously used and published in [39]. The SAM 3D stacks were captured using a time-lapse confocal microscope for every 4 h for 0 to∼80 h and were passed through a pipeline of image correction steps. The mean size of the stacks is 150×512×512 pixels (zyx dimensions).

**Fig 15.**
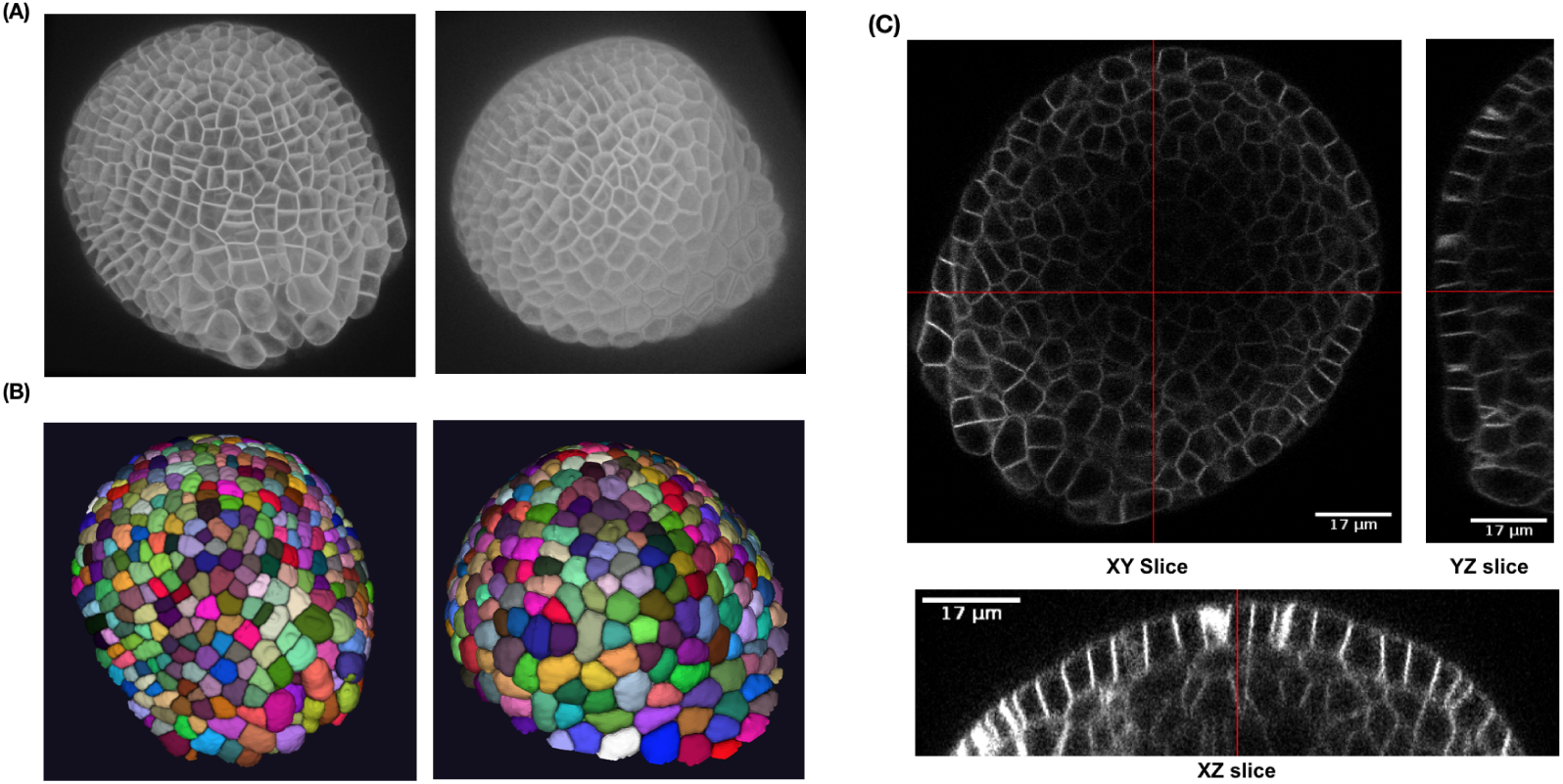
(A) 3D projection of two training images and (B) corresponding ground truth segmentations (C) Lateral (XY) and axial slices (XZ, YZ) of a sample confocal training image

For each stack, slice mis-alignments due to vibrations or microscope stage movements were corrected via translation transformations using the StackReg module of ImageJ. Also, z-slices with horizontal shifts were replaced with the closest z-slice with no shifts. Imaging errors during vertical movement of the plant due to growth were also compensated for by estimating stretching constants for each stack by comparing rapidly acquired low-z-resolution stacks with slowly acquired high-z-resolution stacks [[39],S1,Table 10]. In addition, Gaussian and an alternative-sequential filtering was done for noise removal.

The ground truth data for training (Fig 15B) consists of the 3D segmentations of the above image stacks done by 3D watershed followed by extensive manual corrections on a slice by slice basis. For corrections of segmentation errors, cell boundaries were estimated from the segmented images superposed on the original images for visual inspection. For errors due to over or under-segmentation or missing cells, the noise filter and watershed parameters were adjusted until satisfactory segmentations were obtained for peripheral as well as central zones of the SAM images. In the ground truth images, voxels which belong to the same cell, have the same label.

### Test dataset

For testing the segmentation algorithms, a dataset of ten 3D confocal image stacks was used (described in [48]).Test Set 1 (TS1) contains 6 stacks (0h, 24h, 32h, 72h, 120h, 132h timepoints) are from one meristem (FM1 in [48]). Time points of Test Set 2 (TS2, 26h, 44h, 56h and 69h) correspond to FM6 in Refahi et al. Images of the stacks are shown in Fig 4. These images are from the floral meristem of Arabidopsis from initiation to stage 4. Corrections for alignment and vertical movements are described in [48] . Expert ground truth segmentations of these test stacks were also available to perform numerical comparisons between crops of the segmented results and corresponding ground truths.

To evaluate the segmentations, crop regions were defined on each test stack and corresponding ground truth image. This is done firstly to avoid low intensity regions in the raw images which often correspond to instances of incorrect segmentations in the ground-truth images. Secondly, It was observed that some of the hybrid pipelines faced memory issues when handling very large volume stacks. Therefore, the crops were done such that the volumes of all raw test stacks could be processed by all pipelines. Homogeneous crops of the 10 raw images and corresponding ground truth images were therefore created and used for evaluating all the pipelines for all the experiments described in this work.

### Software and libraries used

The software/libraries used in this work include Timagetk for implementing the MARS algorithm (https://mosaic.gitlabpages.inria.fr/timagetk/index.html), Numpy [49], Matplotlib [50] for plotting the results. Fiji / imagej [51] was used for image visualization. StackReg [52] was used by the creators of the training and test datasets. Tensorflow [53] and Pytorch were used for implementing the deep learning models.

### Training and segmentation details for deep learning pipelines

The four DL algorithms were first trained using the common meristem dataset. The details on how these pipelines could be reproduced are described in the Gitlab repository (Supporting information file S1). For training the deep learning networks and testing, a CUDA enabled NVIDIA Quadro P5000 GPU was used with a Intel Xeon 3 GHz processor. Python 3.x is the default programming language for training/testing all the segmentation pipelines and implementing the evaluation metrics.

#### Plantseg

The residual 3D UNet as described in [29] was used for training on our data. The Residual 3D UNet is a variant of the 3D U-Net and comprises a contracting encoder and an expanding decoder part but uses residual skip connections in each convolutional block in the U-Net. For final segmentation of our test data, all three of the graph partitioning post-processing strategies (GASP, Mutex, Multicut) were found to provide similar results, so GASP was used to obtain all the final segmentations from Plantseg that are evaluated in this work (Fig 16).

**Fig 16.**
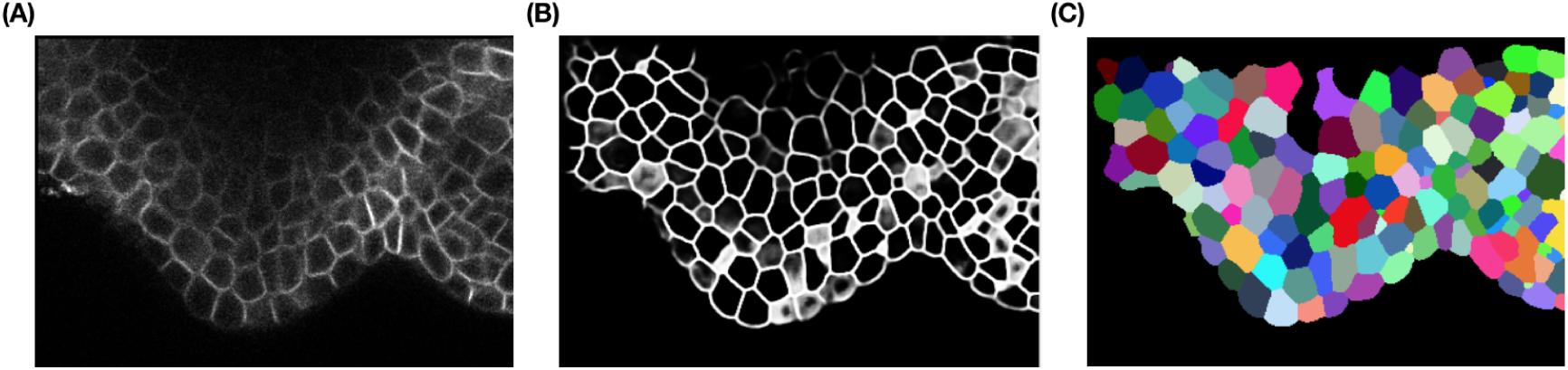
Plantseg workflow: (A) Input image (B) Boundary prediction (C) Final segmentation

The optimal parameters for training the 3D Residual UNet model of the Plantseg pipeline were found through experimentation and the best parameter values are reported here. For training, the Adam optimizer was used with a learning rate of 0.0002. The loss function used was BCEWithLogitsLoss after experimenting with other loss functions like Dice, BCEDiceLoss and BCEWithLogitsLoss. The weight decay was set to 1.0e-05. For data augmentation, random flip and rotation was used along with standardization (Z-score normalization) of the raw image. The number of training epochs required was 500 until the validation loss reduced no further. The evaluation metric used was BoundaryAdaptedRandError. The time taken per epoch is around 550 seconds. Note that Plantseg training configuration allows setting of iterations besides epochs, which was varied between 10000 to 150000 during the experiments.

#### Unet+Watershed

The 3D UNet module of the pipeline was trained using the custom training dataset described above. The 3D Unet predicts 3 classes of output images from an input 3D confocal stack. These classes are: cell centroids, cell membranes and background maps (Fig 17B-D). Dimensions of these 3 output images are the same as that of the input stack.

**Fig 17.**
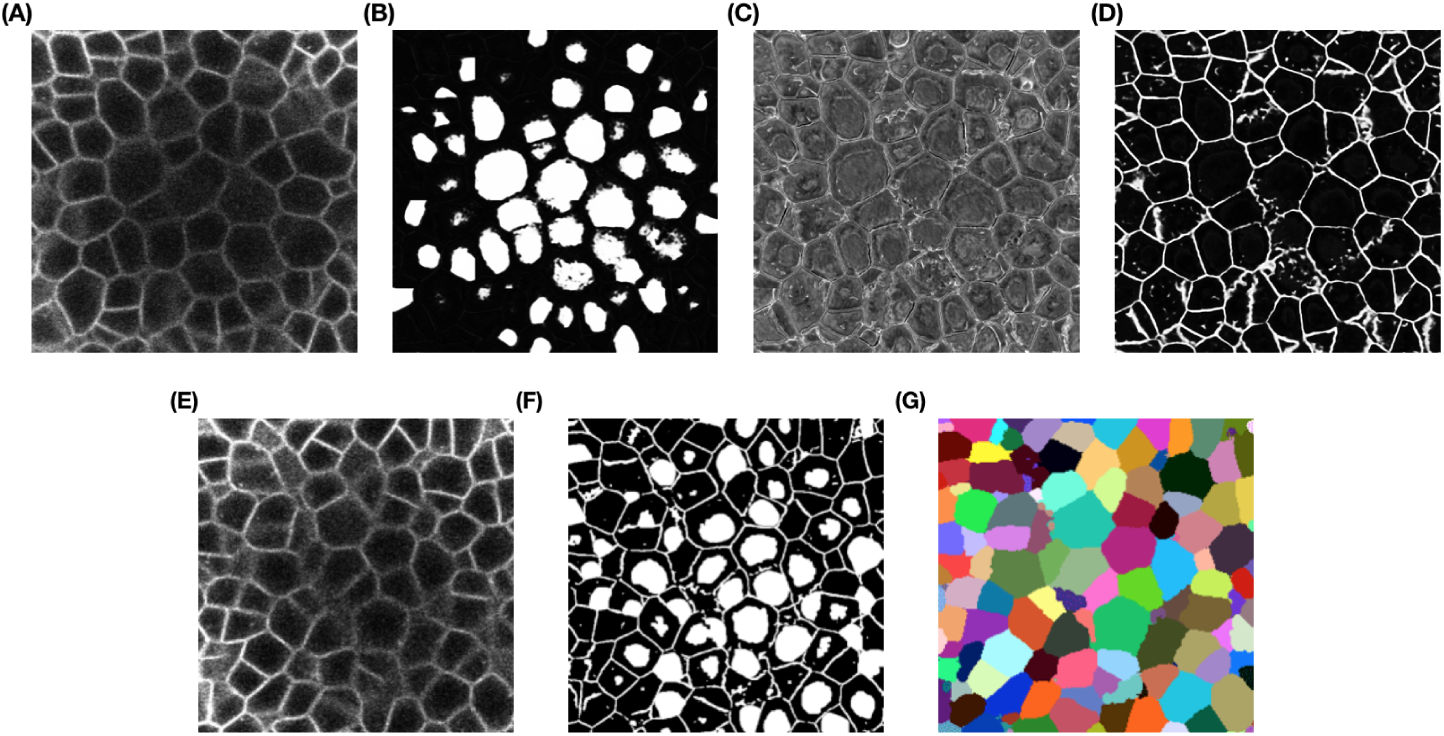
3D UNet+ WS workflow (A) An input confocal image (xy slice) (B) Class 0 prediction-centroids (C) Class 1 prediction- background (D) Class 2 output- cell membranes. (E) An input confocal image (xy slice) (F) Seed image slice (G) Final segmented slice using watershed on (F)

Using these 3 predicted image classes a seed image is obtained by first thresholding the centroid maps (0.8 times the max intensity), followed by its morphological opening (circular kernel, size 5×5×5) and subtracting the membrane and background maps from the resultant image. An example seed image is shown in Fig 17F and the corresponding seeded watershed segmentation output is shown in Fig 17G.

The optimal parameters for the 3D UNet model were found through trials with different numbers of epochs (400, 800), variation of the type of loss function used (mean-squared error and custom loss) and batch size variations of 5 to 20. The optimal number of epochs were found to be 400 and a batch size of 5 was used. The evaluation metric used was mean intersection over union or MIoU. The custom loss function used during training is called weighted_binary_class_crossentropy and is a binary cross entropy based loss with class weighting (0.3 for each of 3 output classes). The optimizer was Adam and a learning rate of 0.001 was used along with batch normalization. For data augmentation, flipping and rotation was used. The UNet watershed pipeline as proposed by the authors does not include any data normalization function in their training configuration. Batch size is set to the default value of 5. The time taken per epoch is around 750 seconds.

#### Cellpose

The Cellpose pipeline uses a 2D UNet to predict horizontal (X) and vertical (Y) flows along with a pixel probability map for each 2D test image. Using the XY intensity gradients or vector fields, pixels belonging to each object to be segmented can be aggregated around the centroid region for that object. For the final segmentation in 3D of the test sets, Cellpose uses the 2D trained model to predict the horizontal and vertical gradients for each of the XY, XZ and YZ sections of a 3D volume. These six predicted gradients are then averaged pairwise to obtain the final XYZ vector map in 3D.

The 2DUNet module was trained using 2D slices (512×512 pixels) from the custom training dataset along with their corresponding 2D masks. The tunable parameters of the method such as flow_threshold, cell probability threshold and cell diameter were set to the default values of 0.4, 0.0 and 30 pixels respectively.

Optimal values of different model parameters used for the Cellpose training included batch_size=8, diameter=30, learning_rate=0.2, weight_decay=0.00001 and momentum = 0.9. Rotation and flipping were used as augmentations. For data normalization, the authors of the pipeline use a custom function which normalizes images so that 0.0 is 1st percentile and 1.0 is 99th percentile of image intensities. The same function was used by us. The model used a SGD optimizer and a SigmoidBinaryCrossEntropyLoss loss function and the training took 2050 epochs. The learning rate hyper-parameter was tuned for this model between 0.0004 to 0.2 to observe the results. The best results were obtained for a learning rate of 0.0008. The time taken per epoch is around 600 seconds.

#### Mask RCNN

Mask RCNN (also called MRCNN in this paper) uses a backbone network such as Resnet-50 or Resnet-101 to extract image features followed by generating region proposals of objects (to be segmented) using a region proposal network (RPN). These region proposals are refined and fed to a fully convolutional classifier to identify object classes. The final output of MRCNN includes 1) boundary box for each object instance 2) pixel level mask for each object identified 3) class predictions for each object instance 4) confidence score of each prediction.

In the MRCNN with watershed pipeline, using a 2D trained Mask-R CNN model the cell regions in each Z slice of a 3D volume are predicted (Fig 18 C). The predicted Z slices with the identified cell regions are stacked together to produce a 3D binary seed image which is then labeled using a 26-neighbor connected components labeling method . The labeled seed image is then used for watershed based post-processing to obtain the final 3D instance segmentation. The Mask-RCNN algorithm with a Resnet 101 backbone network was trained using 2D images along with instance masks for each object (to be segmented) in the image. Using Resnet 50 as a backbone network did not give satisfactory results. Training images (2D slices) from the custom training dataset were used and the instance masks were generated from the corresponding ground truth 2D masks (each labeled cell region forms an instance mask) as shown in Fig 18A. Both the raw image and the set of instance masks for each image are then used for training the MaskRCNN network.

**Fig 18.**
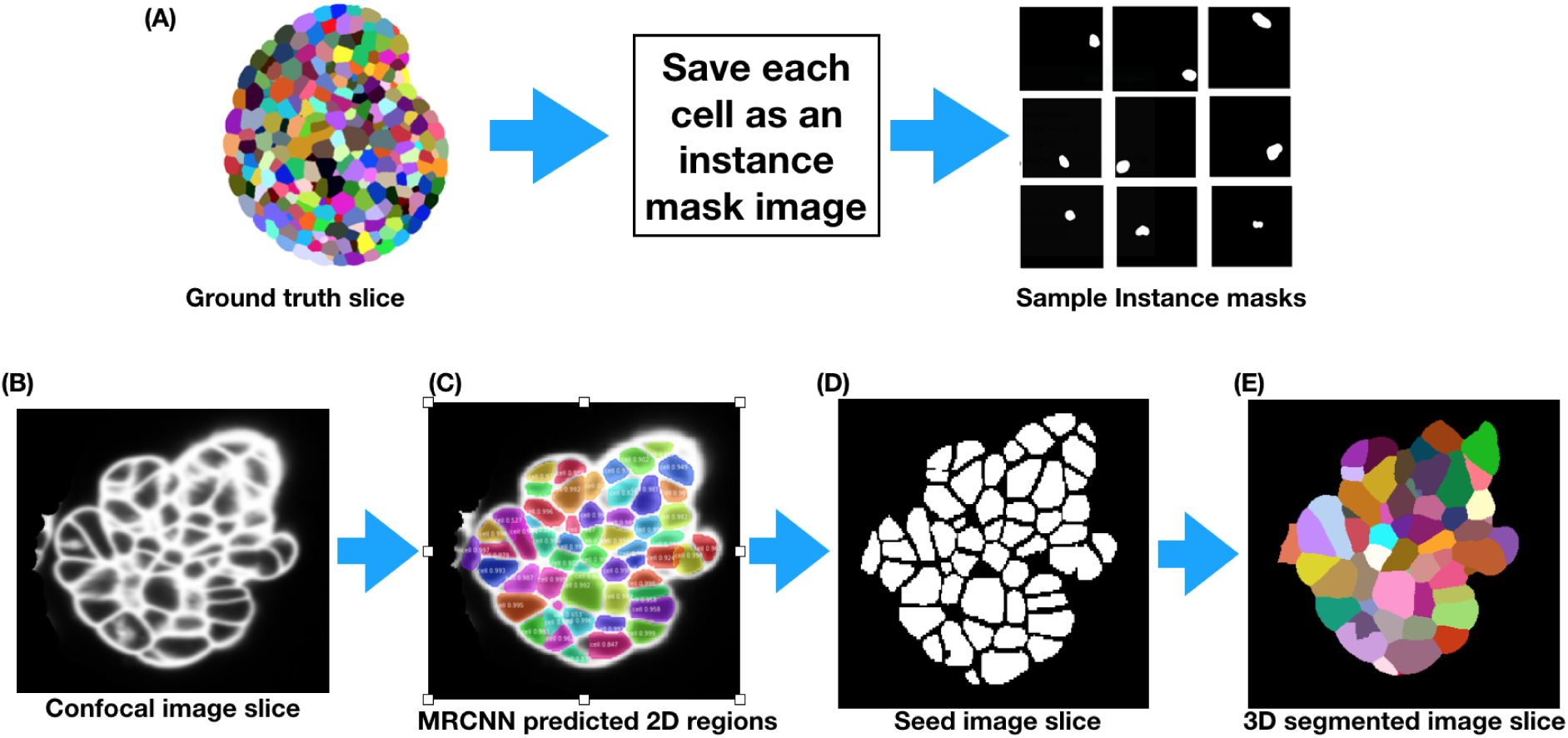
MRCNN+Watershed workflow: (A) Creation of instance masks for training MRCNN (B) Example confocal slice (C) 2D predictions by MRCNN (D) Binary seed image created from identified cell regions in (C).(E) Same slice after 3D segmentation using watershed on the binary seed image.

The MaskRCNN model provides a range of hyper-parameters to tune. For training the MaskRCNN model in this work, the parameters like RPN threshold, RPN anchor scales, number of anchors per image, number of training ROI’s per image etc. were varied and tested through experiments. The optimal values for these were found as follows : RPN threshold =0.9, RPN anchor scales= (8, 16, 32, 64, 128), number of anchors per image, number of training ROI’s per image=300, Detection max instances = 400. An SGD optimizer was used with a learning rate of 0.001, momentum of 0.9 and weight decay of 0.0001. The loss function used was Binary cross entropy. Other loss functions are also defined for this model such as rpn_class_loss (RPN anchor classifier loss), rpn_bbox_loss (RPN bounding box loss), mrcnn_class_loss (loss for the classifier head), mrcnn_bbox_loss (Loss for Mask R-CNN bounding box refinement) values were also monitored during training. The data augmentations used include random flipping and rotation of the images and since no normalization function is defined in the original configuration, we followed the same. The time taken per epoch is around 600 seconds.

The training loss curves obtained at the end of training each of the models from the four deep learning pipelines are plotted in Fig 19. These show the optimal number of training epochs required for the models of the four pipelines (while training on normal images). The training time for the deep learning models in each pipeline are given in Table 2. The parameters to tune for each pipeline is summarized in Table 3.

**Fig 19.**
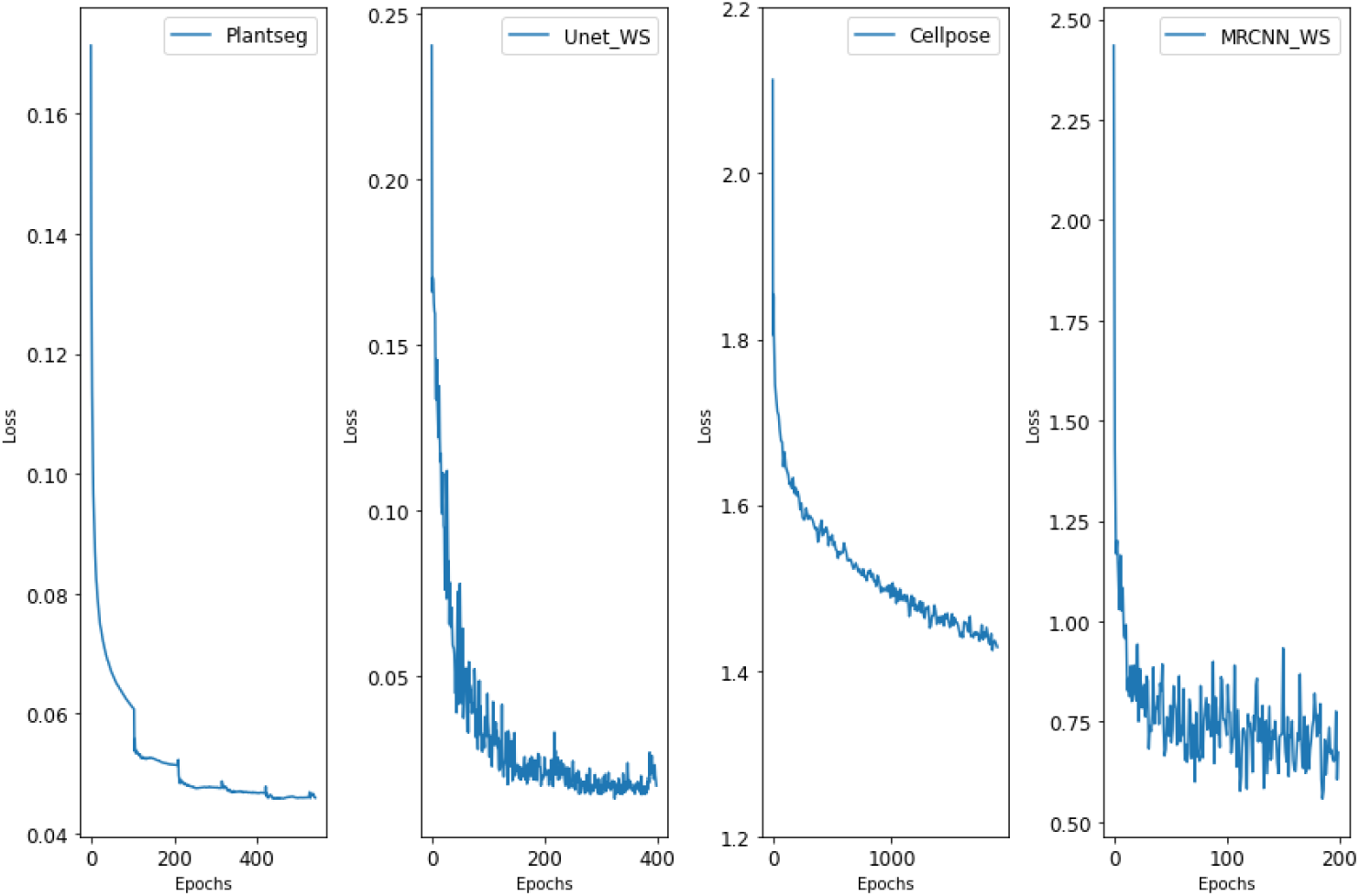
Loss vs epoch plots for training the models from four pipelines

**Table 2.** Training details for all deep learning pipelines

| Pipeline | Deep model | Optimizer | Loss function(s) | Epochs | Training time | Augmentations |
| --- | --- | --- | --- | --- | --- | --- |
| Plantseg | 3D Residual UNet | Adam | BCE with logits loss | 500 | 76 hours | rotate, flip,elastic deformation |
| 3D UNet+WS | 3D UNet | Adam | Weighted binary class cross entropy | 400 | 83 hours | rotate, flip |
| Cellpose | 2D UNet | SGD | Sigmoid Binary Cross Entropy | 2050 | 34 hours | rotate, flip |
| MRCNN+WS | Mask RCNN | SGD | Binary cross entropy | 200 | 33 hours | rotate, flip |

**Table 3:**
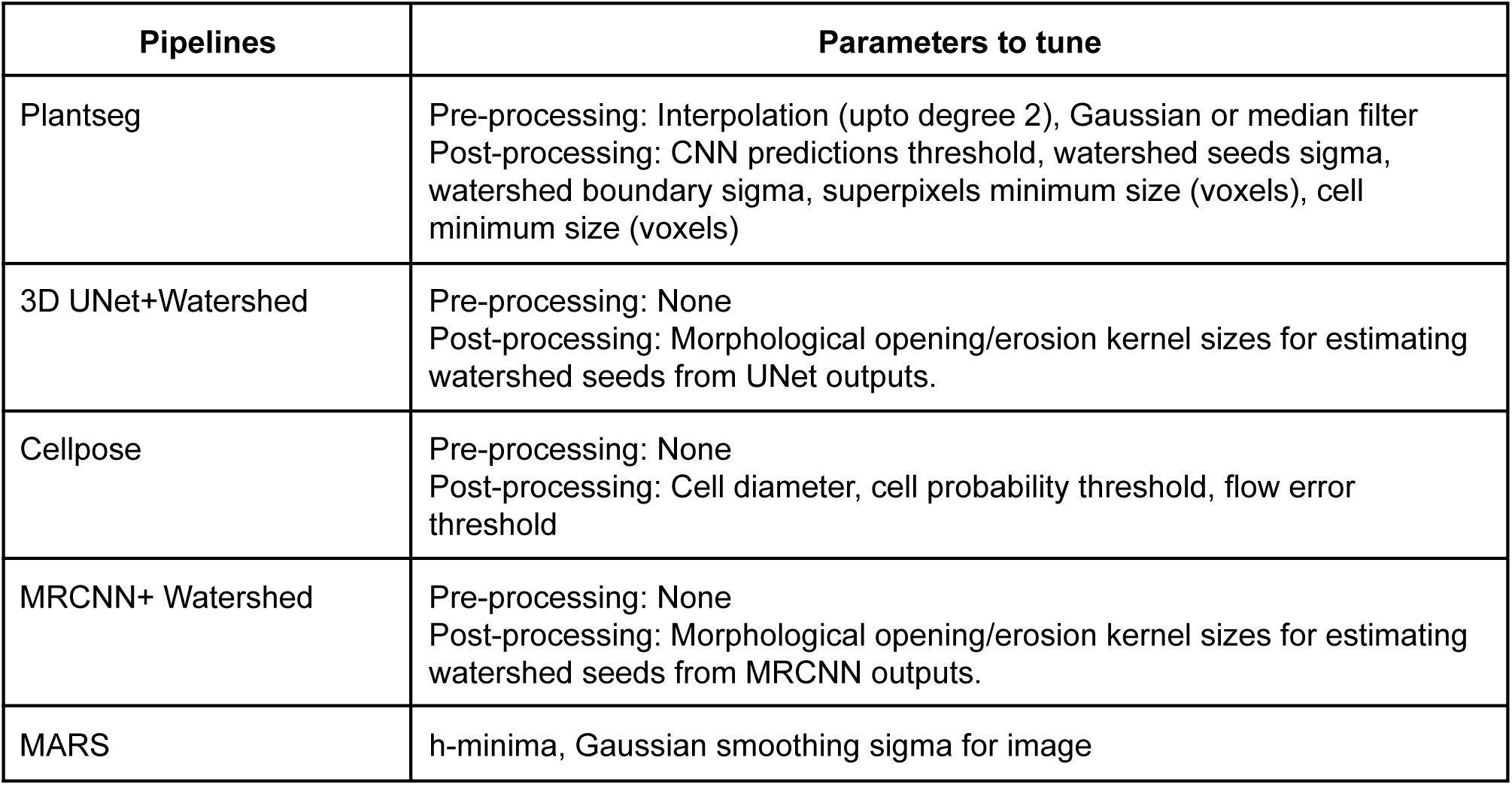
Parameters to tune for each segmentation pipeline

#### MARS

No training is required for MARS but it can require adjusting of its parameters to obtain the best segmentation results. These include the h-minima value and Gaussian smoothing sigma for the images (Table 3). We made an estimate of the execution time of the MARS algorithm on one of our sample test stacks (TS1-132h, volume = 700×700pixels x 297 z slices) and a single run of the algorithm takes 78 seconds on a CPU (measured from the start to finish of the algorithm execution). The parameters to tune include h-minima (integer values starting from 1, upper limit depending on image intensities) and gaussian_sigma (fractional float values varying between 0 and 1) for image smoothing. The initial setting of the parameter values and expertise of the user influences a lot the number of times these parameters need to be adjusted. So for a normal user, trial runs with 10 step changes of gaussian sigma and 10 of the h-minima parameter required a total of around 30 minutes. Additionally, manual supervision is required for inspecting the quality of segmentation in each slice of the stack, which could be estimated to take around 1-2 minutes after every run of the algorithm. Thus the overall time required by MARS for segmenting a large stack like the one used above could reach up to 45 minutes. On the other hand, Plantseg (using a trained model) took 8 minutes for segmenting the same stack on a GPU based computer and did not require any parameter tuning during this phase.

Requirements of parameter tuning for MARS vary from one dataset to another, and may be necessary for stacks within the same dataset. For example MARS parameters used to segment our TS1 and TS2 stacks were hmin= 2 and sigma = 0.4. However, on another dataset (Ovule images, see section Results, Strategy 4: Evaluating pipelines on unseen data types), a lot of tuning was necessary (Fig 20). For this dataset, we had to explore hmin values between 2 to 30 and sigma values in the range 0.2-0.8. Overall more than a 100 combinations of hmin and sigma parameters had to be tested to identify the best one that yields the highest VJI. The best accuracy (given in Fig 12B) was obtained for hmin values of 27, 27 and 9 respectively for the images Ovule 1, 2 and 3 stacks which is very different from the values used for our TS1 and TS2 sets.

**Fig 20.**
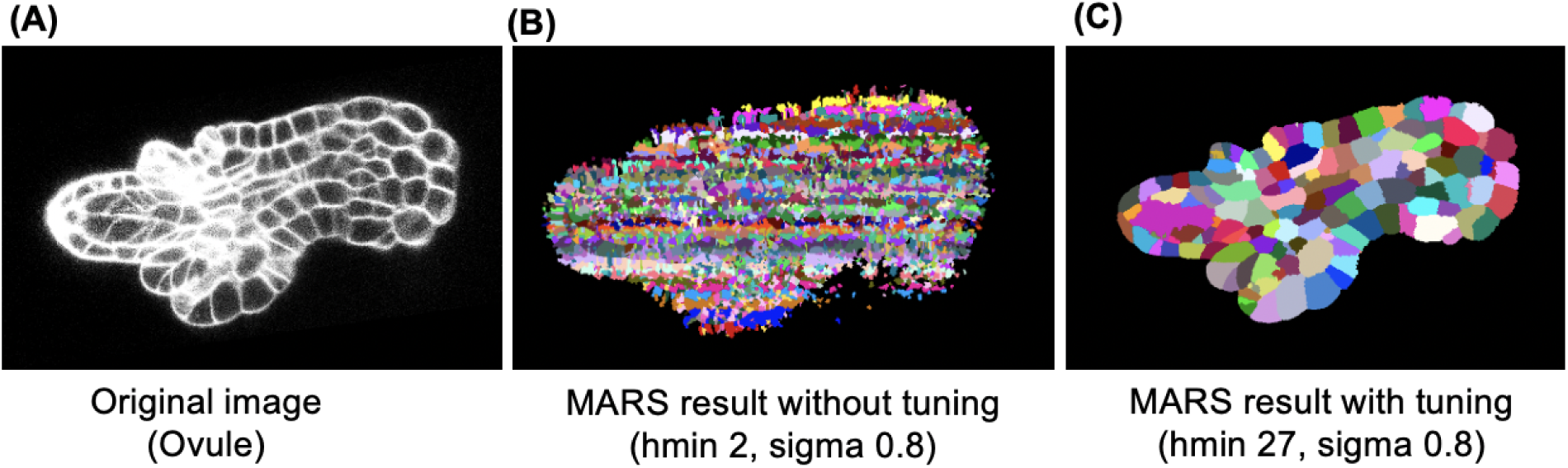
(A) Original ovule image (B) Impact of using hmin= 2 and sigma value =0.8 for MARS (C) Result of MARS on the same image after tuning parameters.

### Evaluation metrics

#### Volume averaged Jaccard index

The Jaccard index is a metric which estimates the similarity between two regions of labeled images, G and P, in terms of the intersection between them divided by their union [54]. Let us denote G_i_ and P_j_ these two overlapping regions, their Jaccard index is defined as:

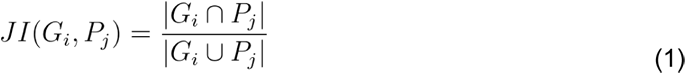

where |X| denotes the volume of region X. We also define an asymmetric inclusion index metric between two regions defined as:

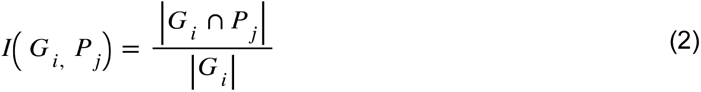

Given a set of *M* labeled regions {G_i_} in the ground truth image G, we associate with each cell region index i a cell region index *A*(*i*) = *j* in the image P containing the *N* predicted segmentations P_j_, using the Jaccard index:

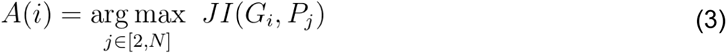

Note that if the cell *G_i_* has no intersection with any cell of P, we set *A*(*i*) = 0. Note that index i varies over [2,M] and j varies over [2,N] as background regions are not included in this estimation and in both the segmented and ground truth images, (the background label is set as 1). Similarly, using the asymmetric index each region *G_i_* of the image G is associated with one region index *B*(*i*) = *j* in the image P according to:

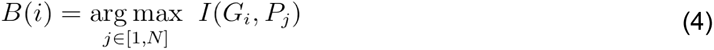

Reciprocally, each region P_j_ of the image P is associated with one region index *B*’(*j*) = *i* in the image G:

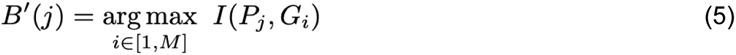

We then define an average metric between two images P and G, called Volume averaged Jaccard, that assesses how well the regions of two images overlap Index (VJI):

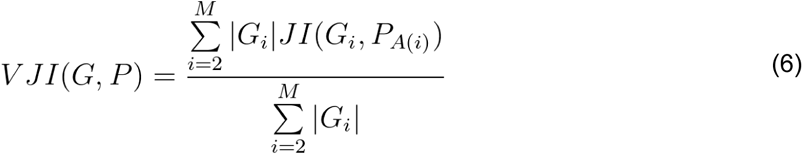

where G_i_ and P_j_ represent regions in respectively images G and P corresponding to either ground truth or predicted labeled cells. In this equation, it is assumed that *JI*(*G_i_, P*_0_) = 0 for any i . Background regions are not included in this estimation and in both the segmented and ground truth images, the background label is set as 1 (index i starts at 2 in (6)).

#### Rates of over and under segmentation to evaluate the quality of 3D segmentation methods

The Volume averaged Jaccard index is a measure that quantifies the degree of overlap between two segmentations although it does not indicate whether the cell segmentation errors are due to over or under-segmentation. In order to detect these different types of errors, we use the asymmetric index (2) to automatically determine a region-to-region correspondence map [55].

Using the previous inclusion index metrics, a correspondence map (Fig 21) is obtained by : 1) making region-to-region association from the segmentation G to segmentation P then 2) repeating the procedure in the opposite direction ie. from image P to image G before 3) building the resulting reciprocal associations between sets of regions.

**Fig 21.**
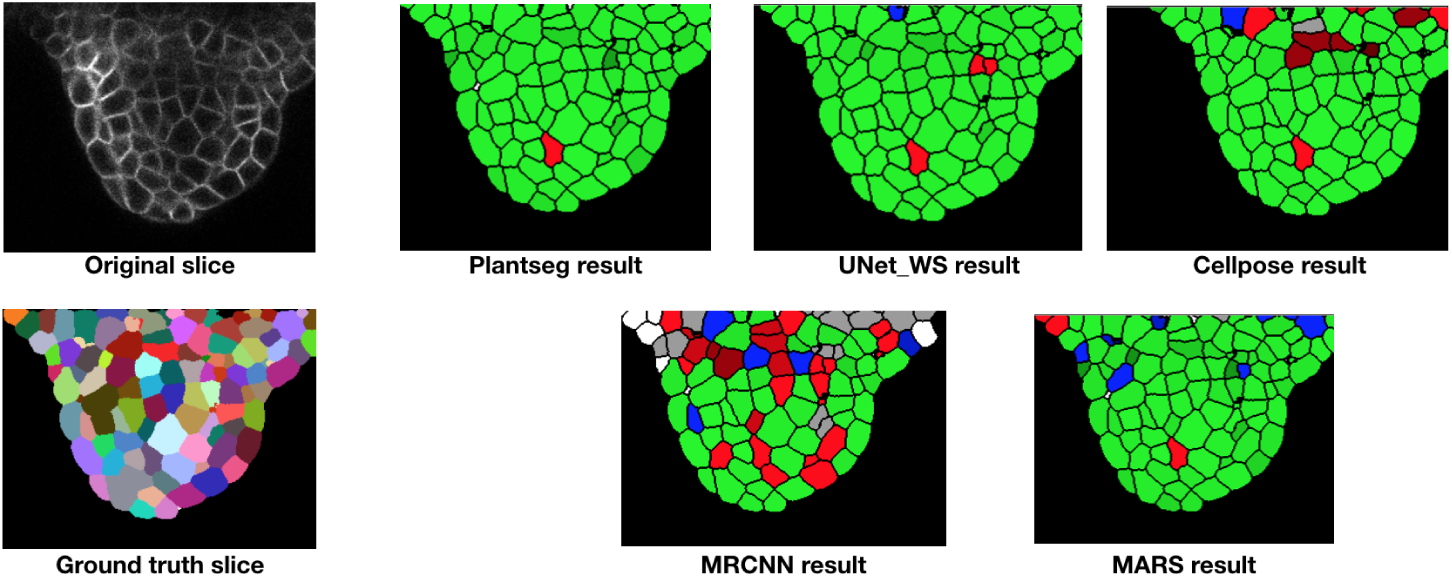
Segmentation quality metric [55] applied to outputs from 5 segmentation pipelines and types of errors displayed as a colormap (on a common Z slice). The green cell regions represent regions of complete overlap between ground truth and predicted segmentations (i.e regions of fully correct segmentation). Red regions represent over and blue regions represent under-segmentation errors. White regions are regions where cells were mistaken for background. The benefit of this metric is that it helps to estimate the rate of over and under-segmentations as a volumetric statistics and as spatial distributions.

The two first steps consist of computing two values *B*(*i*) and *B*’(*j*) for each region index *i* of the first image and each region index *j* of the second image using (4) and (5) respectively. Using the previous computed indexes, set of pair of associated regions index between image P and G can be defined by:

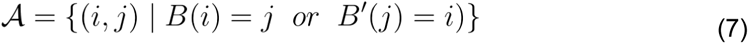

Let us define the two subsets 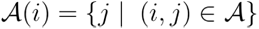 and 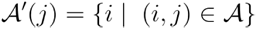 corresponding to the region indexes *j* associated with a given region index *i* and the region indexes *i* associated with a region index *j* respectively. We then consider the different cases of resulting reciprocal mapping:

- one-to-one (exact match between *G_i_* and *P_j_*) if *A*(*i*) = {*j*} *and A’(j) =* {i}.
- one-to-many (over-segmentation of *G_i_*) if 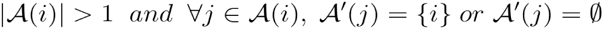.
- many-to-one (under-segmentation of *G_i_*) if 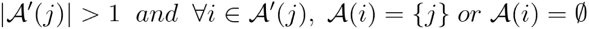.
- many-to-many otherwise

It has to be noted that the correspondence involving the image background is treated separately, ie. Without considering a reciprocal association: the regions of segmentation G that maximize their inclusion with the background of the image P are associated and vice versa. From the resulting reciprocal mapping, a global rate of over and under segmentation can be calculated by counting the number of cells of the over or under correspondence regions in the image P. Because of regions where tissues are not segmented in the ground-truth segmentations, the predicted cells associated with the reference background are not counted in the final rate of over and under segmentation.

### Simulation of image artifacts

The effects of noise, blur and intensity variations are simulated on the test set of 10 confocal images to evaluate their impact on the segmentation quality of the pipelines. The procedure for simulation of these artifacts are described below.

#### Image noise

The Gaussian noise was added to the images. The noise variable z is represented as the probability density function (PDF) P(z) and given by:

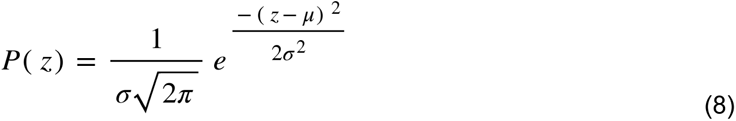

where µ is the mean and σ is the standard deviation (or the square of the variance). Gaussian noise was generated with mean =0.0 and two different values of noise variance ([0.04, 0.08]) and added to an image.

#### Image blur

To simulate motion blur, the test confocal image i(x,y) is convolved with a horizontal motion blur kernel w(dx, dy) of size 4×4 to get the final blurred image b(x,y). The convolution in the spatial domain is defined by the Equation 9 below:

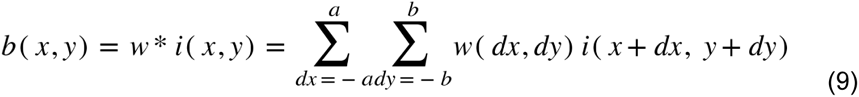

The kernel used for simulating the image blur is a 9×9 horizontal motion blur kernel as shown below.

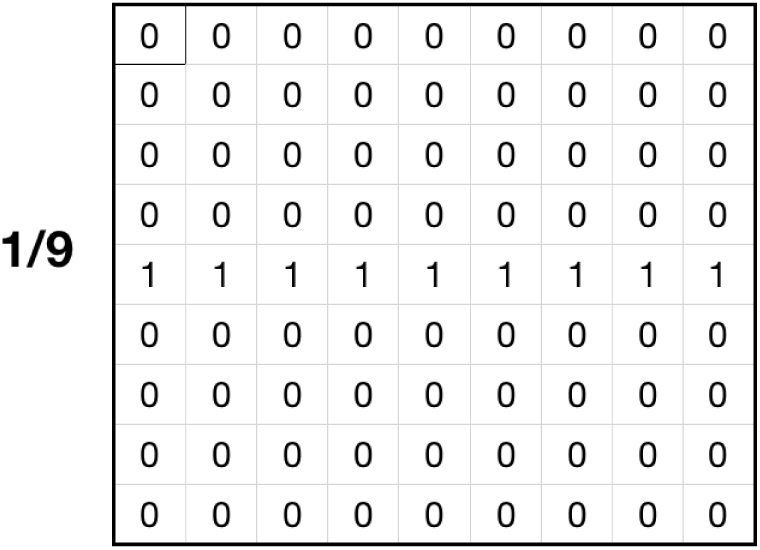

#### Image intensity variations

Partially bright regions in microscopy images may be caused by inhomogeneous illumination sources and shadow effects are mostly caused by presence of light absorbing objects or obstructions [42,43,56,57]. To emulate the effect of intensity variations within an image, partial overexposure (Fig 22A) and random shadow regions (Fig 22B) are imposed (individually) on the test images. In order to impose the partial overexposure effect, for each Z slice of a given 3D test stack, a brightness mask is created, which is a 2D array having the same size as the x, y dimensions of the 3D stack. This 2D brightness mask array is filled with gray integer values of 255 for the left half of the mask array (Fig 22A). This brightness mask is then numerically added to each Z slice array with 30% transparency (using OpenCV function cv2.addweighted) to obtain the partially brightened image array as shown in Fig 22A.

**Fig 22.**
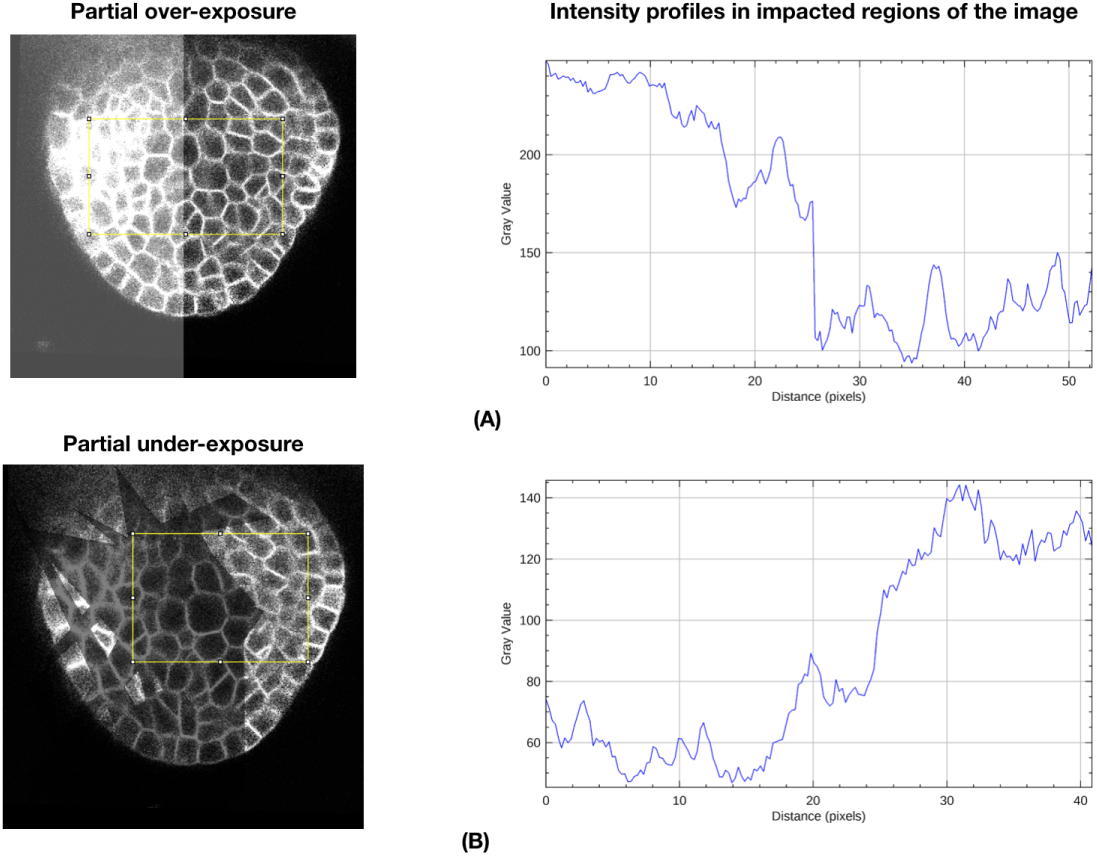
Modification of image intensity (inside selected area within the yellow box) (a) image intensity transition under partial overexposure (b) image intensity variations due to imposition of underexposure.

For creating the randomly shadowed or under-exposed regions, a shadow mask is created with random dark geometrical patch areas for each 2D Z slice of a 3D test stack. The dark patches are created by reducing image pixel intensities within the patch areas by a factor of 30% of the original intensity values. Fig 22 A and 22B shows the intensity profiles within the yellow boundary boxes impacted due to the over and under-exposure effects respectively. Within the over-exposed regions, image intensities are higher than original values and vice-versa for under-exposed regions.

## Acknowledgement

We would like to thank Grégoire Malandain for critical reading of the text, Alexandre Cunha, Johannes Stegmaier for advice on the UNet + WS implementation. Jonathan Legrand is thanked for helping with the implementation of MARS and building the Gitlab repository. We would also like to thank Emmanuel Faure for his guidance on the use of the Morphonet platform.

## Supporting information

## S1 Gitlab repository SegCompare

SegCompare is an open repository of resources to train, test and evaluate multiple deep learning or non-deep learning algorithms (that are described in this article) and compare their segmentation quality both in a quantitative and visual manner. All the segmentation pipelines and evaluation methods in the repository are implemented using Python programming language. SegCompare resources are user friendly and contain:

a. Steps for replicating deep learning and non-deep learning segmentation pipelines on custom user data. This includes steps for retraining the deep learning models used in this article as well as using them for directly segmenting user data.
b. Methods for quantitative evaluation of segmentation quality for a given segmented image and its ground truth image
c. Methods for 3D visualization of segmentation quality on the Morphological browser interface named Morphonet.

Link to Gitlab repository: https://gitlab.inria.fr/mosaic/publications/seg_compare

The Gitlab repository hosts a set of Jupyter notebooks for running the different tools presented in this work-such as implementation of the MARS segmentation pipeline, the segmentation evaluation metrics and also links to sample images with ground truth where the methods can be tested. Link to the notebooks are below: https://gitlab.inria.fr/mosaic/publications/seg_compare/-/tree/master/notebooks

The repository also connects to a documentation page where details about each segmentation pipeline, segmentation evaluation and visualization methods are provided along with instructions for their installation. Further, links to all the datasets used in this work may be also found in this page.

Link to documentation page: https://mosaic.gitlabpages.inria.fr/publications/seg_compare/

**Fig 1.**
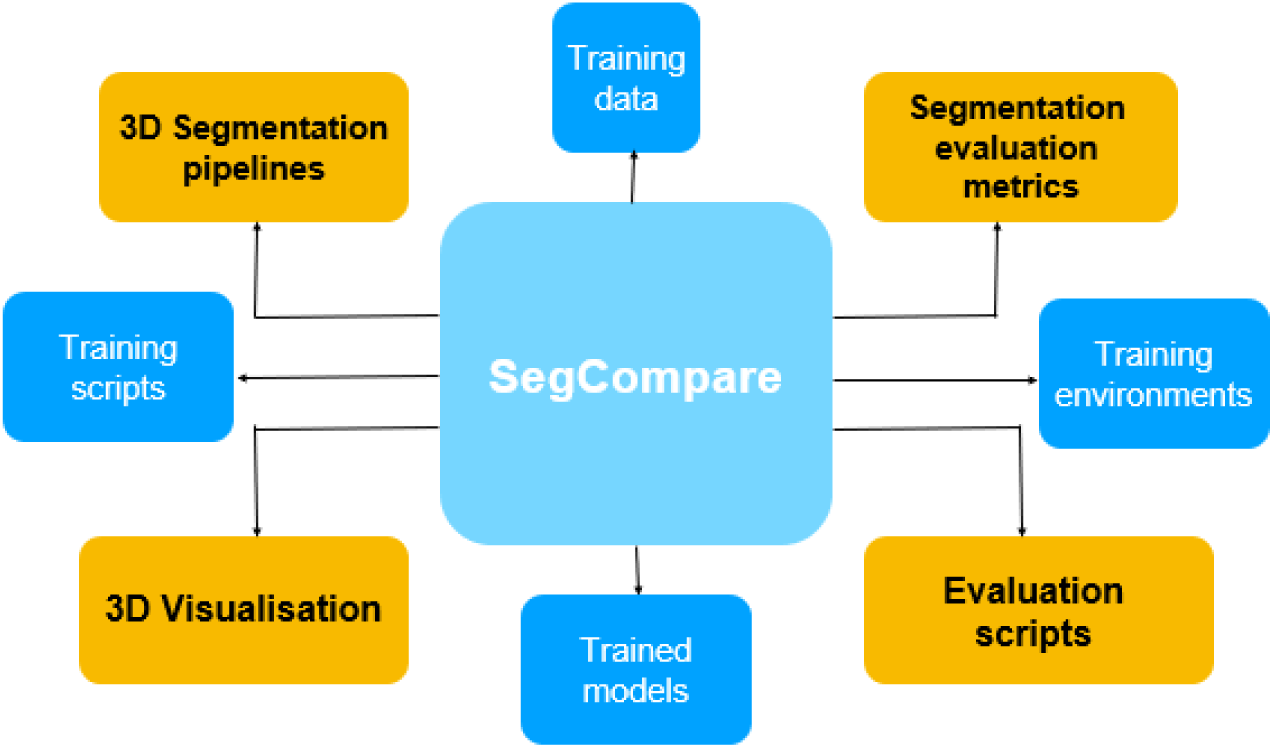
Components of the SegCompare repository on Gitlab hosting the resources for training and evaluation of segmentation pipelines described in this paper.

The utilities of this repository (S1 Fig 1) are the following:

### For users wanting to segment their data

They can directly use one of the trained models and pipelines described here to segment their data.

### For users designing a new 3D segmentation pipeline

They can use the fully annotated image datasets and evaluation methods to estimate the quality of their method.

### For users having segmented data and expert ground truth

They can use this repository to evaluate the quality of their segmentations with quantitative metrics and 3D visualizations.

The contents of SegCompare are as follows:

### Datasets

A description of the confocal image datasets used for training and testing the segmentation pipelines described here. The links to the actual images are provided in Supplementary material S3.

### Installation instructions

Each pipeline has its own library dependencies and therefore needs dedicated environments for running. For this, Python environments for each pipeline may be set up using yaml files, which are also provided in the SegCompare repository. The installation Instructions section provides detailed instructions to install environments for the segmentation pipelines, evaluation and visualization methods. After installing the environments, users can run the segmentation pipelines (for training or testing) or the evaluation and visualization methods.

### Segmentation pipelines

Brief descriptions of each pipeline along with details of their pre- and post processing steps are presented here. Steps for dataset preparation for training the pipeline and links to training dataset are provided.

### Evaluation

This section provides details of the segmentation evaluation metrics (Volume averaged Jaccard Index, Rates of over and under segmentation) and Jupyter (Python) notebooks for implementing them on a pair of segmented images. For evaluating segmentations a user must have a segmented image and corresponding ground truth segmentation (currently .tif format is supported for images). Sample segmented and ground truth data are in the data repository described in Supplementary material S3.

### 3D Visualization

This section describes how the browser based Morphological data visualization platform Morphonet may be used for visualizing segmentation quality. A Jupyter notebook is provided which contains the full implementation of the visualization pipeline starting from a segmented image. For this visualization, results from the Jaccard Index evaluation notebook are required and the full workflow is documented in the repository. Links to sample meshes and datasets for uploading to Morphonet are provided along with demo videos (described in Supplementary Material S3).

### Downloads

This section contains links to Jupyter notebooks for implementations of the MARS pipeline, segmentation evaluation metrics and 3D visualizations. Sample CSV files with Jaccard index estimates and corresponding mesh files for uploading to Morphonet are in the data repository (see Supplementary material S3) .

Link to Jupyter Notebooks on Gitlab: https://gitlab.inria.fr/mosaic/publications/seg_compare/-/tree/master/notebooks

## S2 Morphonet based visualization of segmentation quality

To visualize quality of the five segmentation pipelines on Morphonet, users can click the link below which will directly take them to an uploaded dataset with segmentation quality information from the 5 pipelines (MorphoNet works best with Chrome or Firefox) : https://morphonet.org/icRos2mO

At first only a 3D mesh of the image is displayed. Next, to visualize Jaccard Index info for the pipelines, click on Info-> Click on Info name-> Set colormap. This superposes the VJI values as color-mapped information on the mesh. Multiple meshes and multiple information may be uploaded in this manner.

To upload and visualize information on Morphonet, users need to create an account on Morphonet.org by clicking the “Signup” option on the page or use the guest account provided with this paper. To use this guest account, users may login using : **username:** guest, **password:** guest2021.

To use the Morphonet platform for uploading new data, after logging in to Morphonet, the users may upload meshes for multiple time points and corresponding information to superpose on the meshes. For visualizing segmentation quality on a cell by cell basis, users need to first compute the Volume -averaged Jaccard Index metric using the Volume averaged Jaccard index evaluation.ipynb (may be found under Downloads section of the SegCompare repository), using as input a segmented image and corresponding ground truth segmentation. The output of the Jupyter notebook is a CSV file containing Volume averaged Jaccard Index measure for each cell. This CSV file along with the ground truth segmented image may be used with the 3D_visualization.ipynb notebook (also under Downloads/Notebooks in the SegCompare repository) to do a one step uploading of mesh and numerical information on Morphonet. After running this notebook, users can go to the Morphonet.org page, navigate to their dataset to visualize it. Sample videos demonstrating these operations may be found in our figshare repository as described below under “Videos”.

## S3 Data and model repositories

The training and test datasets used in this work are available online. The training dataset of shoot apical meristems may be found under: https://www.repository.cam.ac.uk/handle/1810/262530

The full set of test images of floral meristems may be found at: https://www.repository.cam.ac.uk/handle/1810/318119

We have used selected images from the above repository. Within this repository, there are overall 6 meristems named FM1-FM6. From this, we selected two meristems FM1 (18 time points) and FM6(5 time points). Out of these, we chose images corresponding to 6 time points from FM1 and 4 from FM6 to create our test dataset. In our test dataset, we renamed the images from FM1 as TS1-00h, TS1-24h, TS1-32h, TS1-72h, TS1-120h, TS1-132h according to the value of the timepoints of the images. The term TS1 represents “Test set 1” . Similarly the images from FM6 are named as TS2-26h, TS2-44h, TS2-56h, TS2-69h where TS2 represents “Test set 2” and the numbers correspond to the timepoints.

A repository of materials generated as part of this study may be found at LINK: https://figshare.com/projects/3D_segmentation_and_evaluation/101120

This repository (3D Segmentation and evaluation) contains the trained models for each of the pipelines, meshes for uploading to Morphonet and corresponding segmentation accuracy files in CSV format. The test images used in this work (TS1-00h, TS1-24h, TS1-32h, TS1-72h, TS1-120h, TS1-132h, TS2-26h, TS2-44h, TS2-56h, TS2-69h) may be found as .tif images under “Test dataset” within this repository.

(Link https://figshare.com/articles/dataset/Test_dataset/16602323)

The overall contents of the repository are the following:

### Trained deep learning models

The models trained in the four deep learning pipelines are provided. Instructions for running them are in the Gitlab repository (Supplementary Material 1).

### Original stacks and segmented data

Segmented confocal stacks by each of the five pipelines are provided along with ground truth stacks for each. Users may test the segmentation evaluation methods using them. Details of using the evaluation function are in the Gitlab (Supplementary Material 1).

### Meshes for Morphonet

Example meshes that might be uploaded to Morphonet are included. Users may test the Morphonet visualization using these and the cellwise VJI values (saved in CSV files). Procedure for the visualization is provided in the Gitlab repository.

### Accuracy results

Cellwise VJI values saved in CSV files are provided for each pipeline. These may be used for projection on Morphonet for 3D visualization of segmentation quality.

### Videos

Videos (.mp4 format) showing examples on how to use the Morphonet based 3D visualization method on a sample test image, videos showing sample training and test data.

### Test dataset

The test images used in this work (TS1-00h, TS1-24h, TS1-32h, TS1-72h, TS1-120h, TS1-132h, TS2-26h, TS2-44h, TS2-56h, TS2-69h)

## S4 Current research on deep learning based instance segmentation techniques

In complement to the survey given in the introduction, we provide here a more extensive overview of the existing deep learning based segmentation methods identifying the major trends of research in this rapidly evolving field. The focus of the survey is on methods that are developed for instance segmentation of images and papers for non-image datasets are excluded. We identified different categories of pipelines, which have been developed to address specific challenges. The research works belonging to each category are discussed below. The main purpose is to illustrate the existing diversity, rather than giving all the details of the individual methods, which is out of the scope of this article.

### Pipelines for end to end 3D instance segmentation

As discussed in the introduction of the main text, end to end 3D (3D input, 3D output) segmentation pipelines have been implemented using either UNet, residual UNet or MaskRCNN architectures. For more details on the UNet and Residual UNet architectures see [1] [2] and [3] for MaskRCNN. Besides the pipelines used here (Plantseg [4], UNet+WS [5] and Cellpose [6]), ([7]) proposed a method which uses 3D images of A. Thaliana and time lapse images of leaf epidermal tissue for training a 3D UNet. This UNet extracts cell boundaries that are processed using 3D watershed along with conditional random fields (a prediction concept which uses contextual information from previous labels).

### Deep learning algorithms for 2D instance segmentation

Mask RCNN is widely used for highly accurate 2D instance segmentation. [8] tested two deep learning architectures for 2D nucleus segmentation i.e. a feature pyramid network (FPN) and a Mask RCNN. This study indicates that Mask RCNN gave superior results. [9] used a modification of the basic MRCNN to perform multi-organ segmentation of human esophageal cancer CT images and mitigate effects of fuzzy organ boundaries and diverse organ shapes in the images. The additional features in the proposed algorithm include a pre-background classification step to improve boundary predictions and use of a custom loss function. In [10] a modified Mask RCNN architecture termed as Panoptic Domain Adaptive Mask R-CNN is developed to achieve unsupervised segmentation of nuclei from histopathology images.

UNets are also used for 2D image segmentation. In [11] a UNet based module followed by post processing steps of thresholding and watershed is used to predict locations of the cells and their nuclei. Multiple deep learning architectures based on UNet, modified UNet and Mask RCNN are tested for 2D nuclear image segmentation in [12] and the Mask RCNN architecture was found to outperform the UNet based models in the 2D segmentation task. It may be noted that for evaluation [8] use F1-score, while [12] use under/oversegmentation and aggregated Jaccard index.

### Deep learning based segmentation pipelines for specific purposes in bioimaging

a. **Deep learning for cell segmentation and tracking:** Deep learning pipelines for instance segmentation coupled with cell tracking have been proposed in several works, such as [13] where the pipeline makes predictions for every cell instance in videos, as well produces temporally connected instance segmentations. Cell instance segmentation in calcium imaging videos is described in [14] which uses temporal information to estimate pixel-wise correlation and shape information to identify cells and classify active and non-active cells. [15] also proposes a modified UNet based model which can be used for tracking cells while dealing with challenging conditions such as crowded cell regions, poor image quality and on data with missing annotations. The method is tested on cell images from mouse muscle stem cells, HeLa cells, and images from developing embryos. Other approaches for implementation of cell segmentation and tracking include [16] and [17]. The latter uses two UNet models to create an architecture named DELTA to first segment the cells followed by tracking lineage reconstruction from time lapse videos of E. coli cells in fluidic medium.
b. **Pipelines for addressing sparse annotations and small training datasets:** For training of deep learning based segmentation models, annotated ground truth data is essential. However, expert annotation of biomedical images (especially 3D datasets) is a highly labor intensive and time consuming process. For this reason, several deep learning pipelines have been developed which can work with sparsely annotated data. These include the method described by [18] which uses only a few fully annotated voxel instances to segment a full 3D stack. In [19] it is demonstrated how varying the contrasts of cell boundaries and a new loss function (weighted cross entropy) could be useful to obtain high accuracy segmentations when a 3D UNet model trained with a small and sparsely annotated training dataset. The issue of sparse annotations is also addressed in works like [20] and [21].
c. **Pipelines for segmenting images with densely packed cells/tissues :** Cell instance segmentation in images where cells appear in dense clusters or in overlapping manner is a common research problem. It is quite challenging as there are high chances of errors in separating each cell. Specially designed deep learning models for segmenting densely packed cell regions are reported in works like [22]. It uses an object detection module called a feature pyramid network, which apparently outperforms MRCNN in this task. The feature pyramid network extracts information of the same image at different scales, in this case the cells and the subcellular nuclear scale. In [23] a hybrid architecture combining UNet and MRCNN is proposed to address effects of crowded and variable sized objects, named Nuclei Segmentation Toolset or NuSeT for nuclei segmentation. The U-Net here is used for semantic level segmentation, the modified MRCNN predicts the instance bounding boxes based on the UNet outputs which are then finally used as seeds for watershed segmentation. [24] uses a combination of two CNNs, that provide a semantic segmentation of nuclear material, followed by a final instance level segmentation of the individual nuclei. The research in [25] performs 3 class classifications on a human tissue image dataset to distinguish between cell boundaries, inside and outside of dense nuclei regions. A new CNN architecture named HoVer-Net is presented in [26] for instance level segmentation of nuclei from histological images where they appear overlapped with each other. This method can further classify the type of nuclei, e.g.the type of cells-such as between tumor and lymphocyte cells from which the nuclei are obtained. Another method specially designed for dense cell clusters in images is [27] which uses two UNet based deep models to predict pixels belonging to the cell regions. Using this output, a final watershed based segmentation is implemented.
d. **Pipelines for segmenting special cell shapes:** In many biological datasets, cells or tissues could have morphologically complex, e.g. very thin or elongated shapes that are difficult to segment by generic pipelines built for regular spherical cell shapes. Deep learning pipelines, with specific DL architectures for segmenting this kind of data have been developed. This includes the method described by [28] which can perform precise segmentation of neural cells which have unconventional structures while also countering challenges like cell division and unclear cell boundaries. An approach combining object detection and segmentation is described in [29] which is successful in high precision detection of small scale and narrow structures of neural cells. Other works include [30] for the segmentation of glial cells, a deep learning model named DeepEM3D-Net in [31] for segmenting 3D neurite images and [32] for segmentation of diversely shaped human organs in images.

### Conclusions

From this literature survey, the following aspects are observed 1) UNet and Mask RCNN are two most common deep learning architectures that are currently used for instance segmentation of biological images 2) The number of works on end to end 3D deep learning for instance segmentation is much lower than that for 2D image datasets. 3) the existing segmentation pipelines have been trained on a wide variety of datasets (plant and animal tissue, cell and nuclei; 2D and 3D still images, videos, time-lapse images) and therefore it is not possible to determine their relative performance levels. 4) For many of the methods surveyed it is not possible to reproduce the pipeline as they are not open source or do not allow retraining. 5) There exists a shortage of large 3D annotated image datasets on plant and animal tissues which are publicly available.

## S5 Effect of retraining deep learning models with artifacts as data augmentation

It is observed from the results in section “***Strategy 3: Evaluating pipelines on synthetically modified images***” that the accuracies of deep learning based segmentation models deteriorate when they are subjected to images with previously unseen artifacts such as over and under-exposure, noise and blur. Out of these, the effects of partial under and overexposure are seen to significantly impact the Plantseg and UNet watershed pipelines. This may be observed from the plots of Fig 7-9 in the main text.

It may however be noted that the deep learning models respective to these two pipelines (i.e. 3D Residual UNet of Plantseg and 3D UNet model of UNet+Watershed) were initially trained on image datasets that did not contain the partial over or under exposure artifacts in them. Therefore the question arises as to what happens when the training set is augmented with images that contain the overexposure and/or the underexposure artifact? For studying this, a new set of experiments were performed by retraining the two deep learning models of these pipelines. In the new experiments the models were retrained by augmenting the training dataset with images where the under and overexposure effects are introduced. Subsequently, the effect of inclusion of artifacts (as augmentations) in training data are investigated. The details of these experiments for each pipeline (Plantseg, UNet+Watershed) are provided below along with the results.

### Experiment 1: Retraining 3D residual UNet of Plantseg by augmenting with over-exposure

In this experiment, the training dataset for the 3D Residual UNet model of Plantseg was augmented with images where the partial overexposure effect is introduced. This augmented training set had normal training stacks as well as stacks with overexposure effects. The function to create the over-exposure was the same as described in this paper under section “Simulation of image artifacts” under “Image intensity variations”.

The results from this experiment are shown in S5 Fig 1 below. The plot shows the comparison between results obtained with the original model alongside those from the retrained model. The retrained model (Aug_Over) was used to segment 3 types of images which include-normal images, images with over-exposure and images with under-exposure. It is observed that as a result of training the model with this augmentation, the segmentation accuracy of the model increases significantly when it is used to segment images with over-exposure as well as normal images. However, the accuracy of segmenting images with under-exposure using this newly trained model is lower than that of the original model. Thus It may be concluded that this model learns to segment both normal and over-exposed data with high accuracy but does not produce good results on under-exposed images.

### Experiment 2: Retraining 3D residual UNet of Plantseg by augmenting with under-exposure

In this experiment, the training dataset for the 3D Residual UNet model was augmented with images where the partial under-exposure effect is introduced. This augmented training set had both normal training stacks and stacks with under-exposure effect. The function to create the under-exposure was the same as described in Section “Simulation of image artifacts” above. The retrained model (Aug_Under) was then used to segment test data to evaluate its performance. For testing, the same test dataset was used-that is normal 10 test stacks as shown in Fig 4A (main text) and the same 10 stacks simulated with over-exposure and under-exposure (i.e 30 test stacks).

The results from this experiment are shown in S5 Fig 1 below. The plot shows the comparison between results obtained with the original model alongside those from the retrained model. As a result of this training data augmentation, the segmentation accuracy of the model increases for under-exposed test data and it also segments normal images with as high accuracy as the original model. The improvement in accuracy for over-exposed images is not as high as the model obtained in Experiment 1. It may be concluded that this model learns to segment both normal and under-exposed data with high accuracy and also produces good results on over-exposed images.

### Experiment 3: Retraining 3D residual UNet of Plantseg pipeline by augmenting with both over and under-exposure

In this experiment, the training dataset for the 3D Residual UNet model was augmented with both over and under-exposed images. This augmented training set therefore had normal training stacks, stacks with over-exposure and stacks with under exposure effect. As before, the functions to create the over and under-exposure were the same as described in Section “Simulation of image artifacts” above. For testing, also three types of images were used-that is normal 10 test stacks as shown in Fig 4A of main text and the same 10 stacks simulated with over-exposure and under-exposure (i.e total of 30 stacks).

The results from this experiment are shown in S5 Fig 1. The plot shows the comparison between results obtained with the original model with those from the retrained model. This model produces high accuracy for all three types of images-i.e normal, over and underexposed ones. The mixed augmentation thus provides an overall improvement in the segmentation quality of the residual 3D UNet model of the Plantseg pipeline.

**Fig 1.**
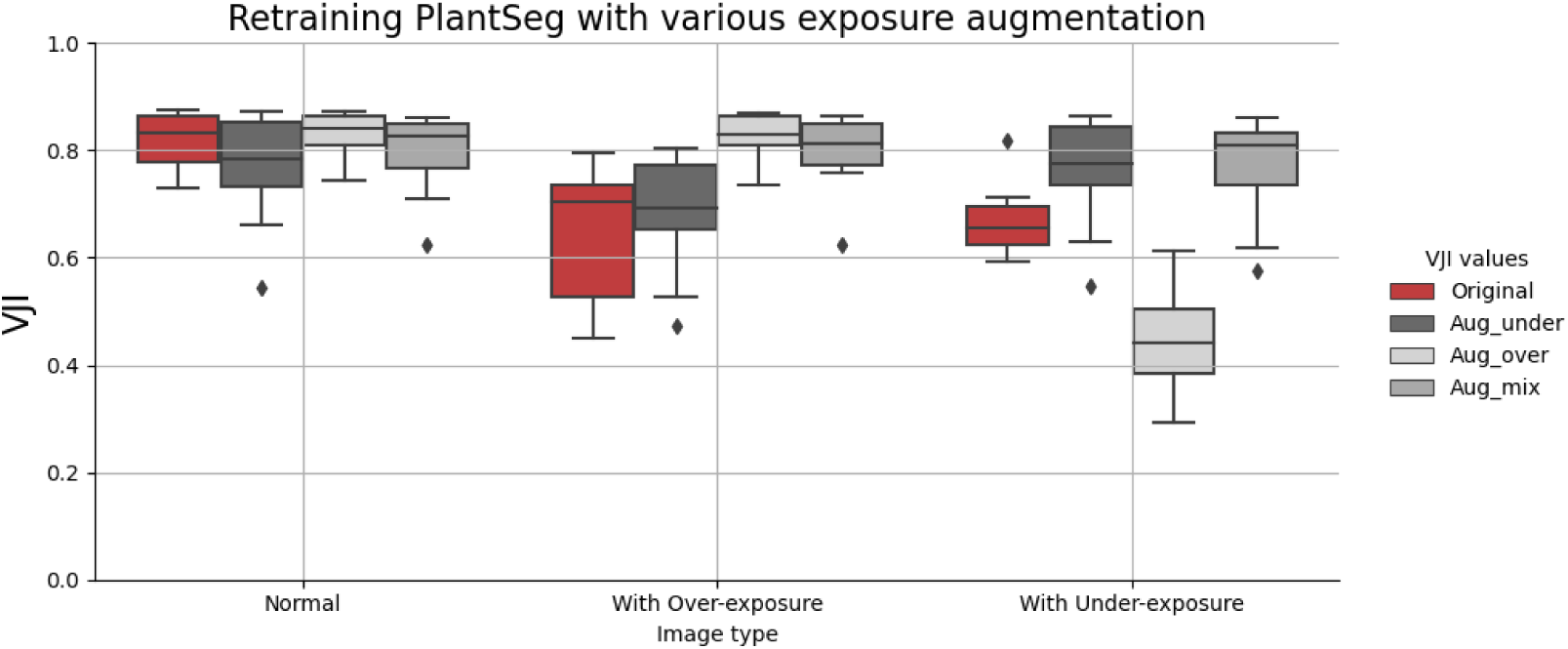
Effect of retraining the residual 3D UNet model from Plantseg on datasets with augmentations. (4 models in total: Original: model trained on unmodified images, Aug_under: model retrained with under-exposed images, Aug_over: model retrained with overexposed images, Aug_mix: model retrained on dataset with over and under-exposed images).

### Experiment 4: Retraining 3D UNet of UNet +WS pipeline by augmenting with overexposure

In this experiment, the training dataset for the 3D UNet model of the UNet +WS was augmented with images where the partial overexposure effect is introduced in the same way as done in Experiment 1 for Plantseg described above. The results from this experiment are shown in S5 Fig 2 below. The plot shows the comparison between results obtained with the original model alongside those from the retrained model. The retrained model (Aug_Over) was used to segment 3 types of images which include-normal images, images with over-exposure and images with under-exposure. It is observed that as a result of training the model with this augmentation, the segmentation accuracy of the model only increases when it is used to segment images with over-exposure. However, the accuracy level of this model for segmenting normal images is much lower compared to the original model. Similarly, accuracy of the retrained model is lower than the original model when trying to segment under exposed images.

Thus it is observed that with over exposure augmentation, the retrained model learns to better segment overexposed images but fails to generalize while segmenting both normal and under-exposed images. This is different from what is observed with the Plantseg pipeline above.

### Experiment 5: Retraining 3D UNet of UNet +WS pipeline by augmenting with underexposure

In this experiment, the training dataset for the 3D UNet model was augmented with images influenced with the partial under-exposure effect. This augmented training set had both normal training stacks and stacks with under-exposure (same as done in Experiment 2 above). The retrained model (Aug_Under) was then used to segment test data to evaluate its performance. For testing, the same test dataset of 30 stacks was used as done in the experiments above-that is 10 normal test stacks, same 10 stacks simulated with over-exposure and under-exposure.

**Fig 2.**
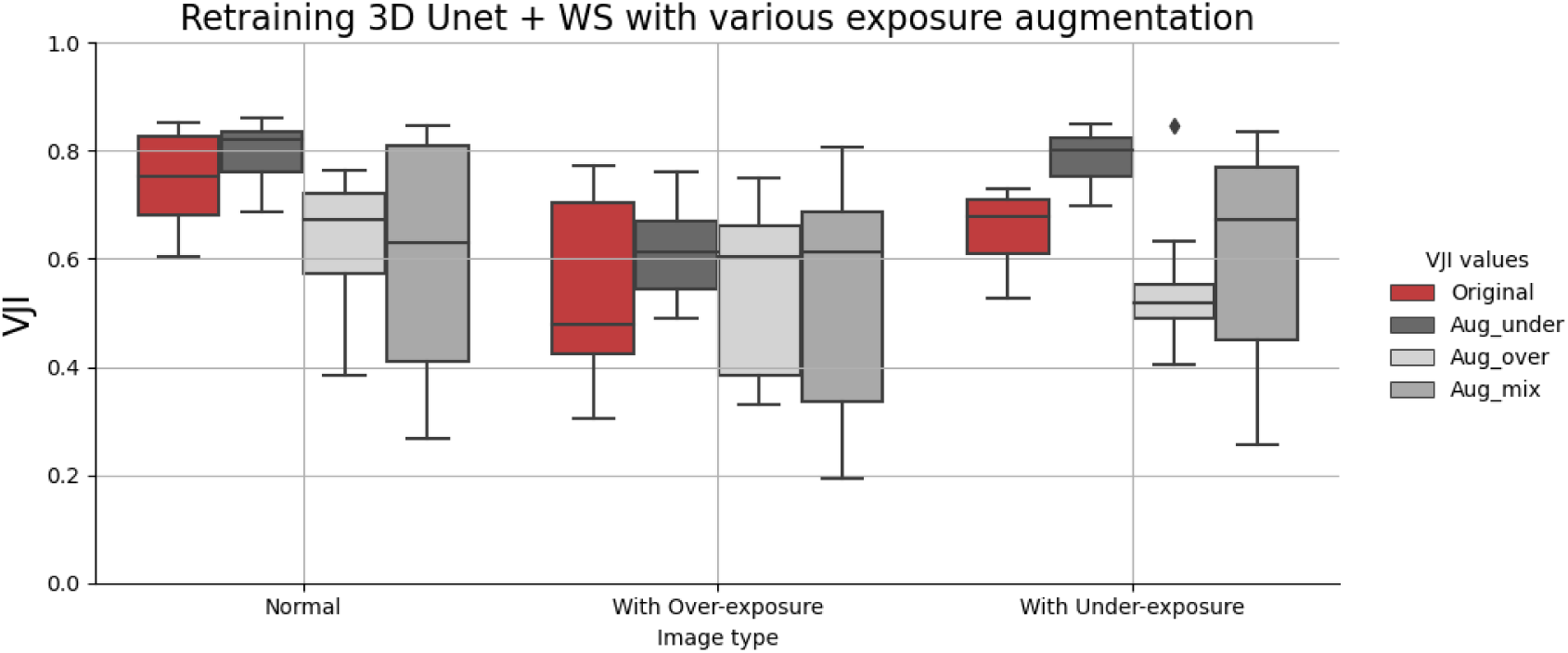
Results from retraining the 3D UNet model from UNet+WS pipeline on datasets with augmentations (4 models in total: Original: 3D UNet model trained on unmodified images, Aug_under: 3D UNet model retrained with under-exposed images, Aug_over: 3D UNet model retrained with overexposed images, Aug_mix: 3D UNet model retrained on dataset containing both over and under-exposed images).

The results from this experiment are in S5 Fig 2 which shows the comparison between results obtained with the original model beside those from the retrained model on augmented data. It is seen that as a result of augmenting training data with underexposed images, the segmentation accuracy of the model increases for all three types of test images. Thus the retrained model performs better than the original model as it provides higher accuracy on normal images as well as on images with under and overexposure. Thus it may be stated that augmenting with under-exposure improves the generalizability of the model.

### Experiment 6: Retraining 3D UNet of UNet+WS pipeline by augmenting with both over and under-exposure

In this experiment, the training dataset for the 3D UNet model was augmented with both images with partial over and under-exposure effects. Also for testing, the same test set was used as in all the experiments above. The results from this experiment are also shown in S5 Fig 2 above. The plot shows that this augmentation improves the accuracy for the over-exposed images whereas those for normal and underexposed ones remains the same as the original results (i.e from model trained on normal images only).

Thus the augmentation with under-exposed images (Experiment 5) is the most effective one for this model as revealed from the experiments. This augmentation produces an overall improvement in the segmentation quality of the UNet+WS pipeline.

### Observations

The main observation from these experiments is that data augmentations could indeed help in improving the performance of deep learning based segmentation pipelines.The Plantseg pipeline results could be improved overall by adding both over and underexposure artifacts in the training set and the UNet+WS results could be improved using underexposure artifact as training data augmentation. However, it is also seen that different deep learning pipelines behave differently in response to data augmentations. While some pipelines may show improved results upon adding certain artifacts for augmenting the training dataset, another pipeline could be adversely affected by the same augmentation. Thus the best training dataset for each pipeline needs to be determined experimentally by observing their response to different training data augmentation strategies.

